# Modeling pancreatic cancer tumor stroma co-evolution in an *in ovo* model

**DOI:** 10.64898/2026.06.26.734719

**Authors:** Raphela Ranjan, Aashreya Ravichandra, Philipp Putze, Anna Chernysheva, Johannes Wirth, Danielle Lucarelli, Wing Yin Ng, Oleksandra Pavlovska, Khanyile Sibusisiwe Sibanda, Erik Leipe, Felix Schicktanz, Stefanie Bärthel, A. Melissa Schlitter, Rupert Ollinger, Marc Ringelhan, Carlo Maurer, Caroline Mogler, Roman Nawroth, Roland M. Schmid, Günter Schneider, Roland Rad, Katja Steiger, Dieter Saur, Maximilian Reichert

## Abstract

Pancreatic ductal adenocarcinoma (PDAC) is characterized by a dense, desmoplastic microenvironment that drives disease progression, yet conventional models fail to capture this complex tumor-stroma coevolution. Here, we utilize the chick chorioallantoic membrane (CAM) platform to investigate tumor-stroma interactions using murine PDAC cell lines and patient-derived organoids (PDOs). Integrating single-cell RNA sequencing and spatial transcriptomics, we show that the CAM microenvironment supports the emergence of complex tumor ecosystems while preserving patient-specific characteristics. Within five days, *in ovo* tumors faithfully recapitulated the structural and molecular features of parental tumors. Histological analysis revealed the rapid recruitment and spatial organization of heterogeneous host cancer-associated fibroblast (CAF) populations, showcasing distinct myofibroblastic and inflammatory stromal states. Crucially, the model preserved intrinsic tumor heterogeneity and permitted functional interrogation of subtype-specific extracellular matrix remodeling and metastatic dissemination. Together, our findings demonstrate that the CAM provides a highly permissive niche for tumor-stroma coevolution. As a rapid, scalable, and biologically relevant platform, this *in ovo* model offers a powerful approach for studying stromal composition, metastatic progression, and patient-specific tumor biology in pancreatic cancer.

## Introduction

PDAC diagnosis and mortality are on the rise owing to a long asymptomatic phase of the disease that prevents effective surgical and/or therapeutic intervention (Wood et al., 2022). Histological, molecular, and transcriptomic evaluations of PDAC tumors have revealed a high level of intra- and inter-tumor heterogeneity, accompanied by a desmoplastic tumor microenvironment (TME) rich in CAFs and their secreted extracellular matrix (ECM) components (Chen et al., 2021). In recent years, the 5-year survival rate of PDAC has nearly doubled (from 6% to 10%) due to advances in combinatorial and/or novel chemotherapeutic strategies (Halbrook et al., 2023). However, frequent tumor relapse accompanied by aggressive metastatic lesions and chemoresistance to therapy regimens has hindered curative efforts (Zhou et al., 2021).

Multiple *in vitro* and *in vivo* strategies have been employed to understand the molecular and genetic mechanisms surrounding PDAC progression and chemoresistance (Halbrook et al., 2023). The isolation and generation of PDAC tumor cells from human and murine tumor tissue have been important and cost-effective tools for understanding tumor cell-intrinsic mechanisms. However, the phenotypic drift and loss of genetic heterogeneity observed in 2D cells have prompted the development of novel 3D *in vitro* strategies (Proietto et al., 2023). To this end, recent advances in organoid technologies have enabled us to maintain primary tumors’ genetic and molecular characteristics, thereby better recapitulating human PDAC. Indeed, PDOs are a crucial tool in advancing the field of personalized medicine and precision therapy (Bose et al., 2021). High-throughput drug screens and pharmacotyping studies using PDOs have identified drug vulnerabilities, thereby providing guidance for postoperative adjuvant chemotherapy selection in pancreatic cancer (Seppala et al., 2022). However, PDOs inherently lack the stromal components that shape PDAC progression and therapeutic resistance, and current co-culture models capture only limited aspects of the desmoplastic stroma (Sharpe et al., 2024).

To overcome this challenge and study the dynamic progression of PDAC, genetically modified murine models (GEMMs) have been employed. The most widely used of these are the KC (Pdx-1-Cre; K-ras^LSL.*G12D/+*^) and KPC (Pdx-1-Cre; K-ras^LSL.G12D/+^; Trp53^R172H/+^) mouse models (Hingorani et al., 2005). The mutant LSL (lox-stop-lox)-KRAS^G12D^ background has been further exploited to include other genes implicated in PDAC progression, including but not limited to *CDKN2A* and *SMAD4*, that have provided insight into liver and lung metastasis as well as cystic-type lesions, similar to that of human PDAC progression (Miquel et al., 2021). These models have also deconvoluted the highly complex stromal compartment, providing insight into CAF heterogeneity and the immune microenvironment (Ferrara et al., 2024). As such, GEMMs, such as the KC and KPC models, offer powerful tools to investigate PDAC biology within a native stromal context; however, they are resource-intensive and poorly suited for high-throughput or patient-specific applications. In addition, traditional mouse xenografts, which involve the orthotopic or heterotopic transplantation of 2D cell lines, PDOs, or tissue specimens into immunodeficient mice, though more scalable, require lengthy engraftment periods and are barriers to rapid translational feedback (Ehlen et al., 2020; Yu et al., 2021).

The CAM assay has previously been employed as an alternative *in vivo* strategy to study cancer biology, in accordance with the 3R principles that advocate replacing, refining, and reducing the use of animals in research (Ribatti, 2023). Compared to conventional murine models, the CAM is a time-, cost-, and labor-efficient model that has been used to investigate multiple phenotypic and functional characteristics of tumor cells, including growth, invasion, and dissemination (Chu et al., 2022; Miebach et al., 2022). However, CAM has not been thoroughly explored to model the complex biology of PDAC, and its full potential as a functional readout of tumor–stromal co-evolution remains largely untapped. Considering the vast heterogeneity of PDAC, ranging from molecularly distinct subtypes to unique sub- TMEs, both of which impact clinical outcomes, our primary aim was to establish the CAM model to study PDAC biology. To achieve this, we leveraged both the translational potential of PDOs and KPC-derived murine cell lines, yielding novel hybrid model systems, PDO-on-CAM and KPC-on-CAM, respectively. By integrating sc-RNAseq and spatial transcriptomic analyses, we assessed the CAM niche’s ability to restore cellular identities and reconstruct the spatial compartments characteristic of native tumors.

## Methods

### Clinical data

Informed written consent was obtained from all patients used in this study in accordance with the Helsinki Declaration and were approved by the ethics review board of Klinikum rechts der Isar, TUM School of Medicine and Health, Technical University of Munich, was received for all patient samples used in this study (Project 207/15, 1946/07, 330/19, 181/17S and 80/17S). The following clinical data were obtained from the hospital information system: Tumor localization, TNM stage, Grading, and source (Supplementary Table 1).

### Establishment and culturing of PDOs

PDOs were freshly isolated from EUS-guided fine needle aspiration/biopsy and surgical resections, following previously published protocols (Dantes et al., 2020). Briefly, tumor specimens were minced into small pieces and digested in a digestion buffer (DMEM-F12 (Cat. 31330095, Thermo Fisher Scientific) containing 6 mg/ml collagenase II (Cat. 17101015, Thermo Fisher Scientific). Tissue pellets were incubated in ACK Lysing Buffer (Cat. A1049201, Thermo Fisher Scientific) and then further digested in TrypLE (Cat. 12604039, Thermo Fisher Scientific) to create a single-cell suspension. Cell pellets were resuspended in 40 μl Matrigel/well (Cat. 354230, Corning Life Sciences). PDO medium (DMEM-F12 (Cat. 11320033, Thermo Fisher), 5 mg/ml D-Glucose (Cat. G8270, Sigma-Aldrich, St. Louis, USA), 0.5% ITS Premix (Cat. 354350, Corning Life Sciences), 5 nM 3,3,5-triiodo-L-thyronine (Cat. T0821, Sigma-Aldrich), 1 µM dexamethasone (Cat. D175, Sigma-Aldrich), 100 ng/ml cholera toxin (Cat. C9903, Sigma-Aldrich), 1% penicillin/streptomycin (Cat. 15140122, Thermo Fisher Scientific), 5% NU-Serum IV (Cat. 355500, Corning Life Sciences), 25 µg/ml bovine pituitary extract (Cat. P1167, Sigma-Aldrich), 10 mM nicotinamide (Cat. N3376, Sigma-Aldrich), 100 µg/ml primocin (Cat. ant-pm05, Invivogen), 0.5 µm A83-01 (Cat. 2939, Tocris, Bristol, UK), 10% RSPO1-conditioned medium (R-spondin-1 overexpressing cell line HEK293T, provided by the Hubrecht Institute, Uppsalalaan 8, 3584 CT Utrecht, Netherlands), 100 ng/ml recombinant human heregulin-1 (Cat. 100-03, Peprotech, Cranbury, USA), and 10 µM rho kinase inhibitor (Cat. TB1254-GMP, Tocris) was added 30 minutes later. For continuous passaging of PDOs, Matrigel domes were dissociated with cell recovery solution (Cat. 11543560, Corning Life Sciences) and ice-cold Phosphate Buffered Saline (PBS) (Cat. 14190144, Thermo Fisher Scientific). After a 30-minute incubation on ice, the organoids were centrifuged at 1200 RPM for 5 minutes at 4 °C, then resuspended in 40 μL of Matrigel/well. They were then cultured in PDO medium.

### Genomic DNA isolation and whole-exome sequencing (WES)

Genomic DNA was isolated from all PDO lines and reference blood using the AllPrep DNA/RNA Micro Kit (#80284, Qiagen, Hilden, Germany) according to the manufactureŕs protocols. DNA concentration and quality were determined using the Qubit 3.0 system (Thermo Fischer Scientific). Library preparations for WES were performed using the Agilent SureSelectXT Low input exome-seq Human v7 kit. Sequencing was performed on the Illumina NovaSeq 600 sequencer, resulting in approximately 140 million 100bp-long paired-end reads per sample. Alignment and mutation calling were performed according to the GATK Best Practices. Read trimming was performed using Trimmomatic 0.38 (LEADING: 25 TRAILING: 25 MINLEN: 50), followed by alignment to the human reference genome (GRCh38.p7) using BWA-MEM 0.7.17. Post-processing was performed using Picard 2.18.26 and GATK 4.1.0.0 with default settings. Somatic mutations were called using MuTect2 v4.1.0.0 (default settings). Mutations with a minimum of two reads supporting the alternate allele and a base coverage of at least 10 in both tumor and germline were retained. Single-nucleotide variants (SNVs) and insertions/deletions (Indels) Σ10 base pairs) were annotated using SnpEff 4.3t, referencing EMBL 92. Copy number variations were detected using Copywriter 2.6.1.2 with default settings.

### PDO proliferation assay

PDO proliferation assay was performed using the CellTiter-Glo® 3D Cell Viability Assay (Cat. G9681, Promega). Briefly, a single-cell suspension of PDOs was generated using 1X Trypsin-EDTA (Cat. 15400054, Thermo Fisher Scientific), and the cells were counted. To avoid 2D growth, wells were coated with 10 µl of 20% Matrigel diluted in PBS. Next, 1000 cells/well were seeded in 100 µl of 10% Matrigel diluted in PDO media. At stated time points, CellTiter-Glo® reagent was prepared according to the manufacturer’s instructions. To determine viable cells, 25 µL of CellTiter-Glo Reagent was added to each well, resulting in cell lysis and luminescence proportional to the ATP present at the time of lysis, a value directly linked to the number of viable cells. Luminescence was measured every other day for 10 days, and the proliferation rate of the PDO lines at each time point was quantified by normalizing to Day 0 luminescence.

### Generation of 2D cell lines from PDOs

2D cell lines were generated from PDOs by allowing over-confluent organoids to outgrow and attach to the bottom of the well. Upon routine culturing of organoids, the remaining 2D cells were expanded and cultured in RPMI (Cat. 11875093, Thermo Fisher Scientific) supplemented with 10% Fetal Bovine Serum (FBS) (Cat. A5256701, Thermo Fisher Scientific) and 1% Penicillin/Streptomycin cocktail (Cat. 15070063, Thermo Fisher Scientific).

### Murine 2D cell lines

Primary murine PDAC cell lines were isolated and established from tumor-bearing endogenous mouse models and were kindly gifted by Prof. Dr. Dieter Saur. Cell lines 8442, 9591, 53631, 16992, and 9091 were isolated and established from Ptf1a^Cre+/-^; K-ras*^G12D/+^*mice. Cell line 9366 was isolated and established from Pdx1^Cre^ ^+/-^; Kras ^G*12D*/+^; Trp53 ^R172H/+^ mice. Cell lines 12548 and R405 were isolated and established from the mouse model P48^Cre^ ^+/-^; Kras *^G12D^*^/+^; Trp53 ^R172H/+^ with tdTomato and tdTomato-EGFP, respectively. Cell lines R4152 and R3924 were isolated and established from the mouse model Pdx1^Cre^ ^+/-^; Kras *^G12D^*^/+^ with tdTomato. Cells were authenticated via genotyping. Murine cells were cultured under standard sterile conditions in DMEM 4.5 g/L glucose medium (DMEM high glucose, Cat. 11965092, Thermo Fisher Scientific) supplemented with 10% Fetal Bovine Serum (FBS) (Cat. A5256701, Thermo Fisher Scientific) and 1% Penicillin/Streptomycin cocktail (Cat. 15070063, Thermo Fisher Scientific).

### PSC-CAF activation

Human pancreatic stellate cells (hPSCs) were purchased from InnoProt (Cat. P10474). Murine PSCs (mPSCs) were isolated as previously described (Feldmann et al., 2021). Supernatant from PDO-generated 2D cells or murine 2D cells was collected at 80% confluency, filtered through a 40 µm filter, and stored at -20°C. hPSCs or mPSCs were cultured for 1 passage in HSC-supplemented medium (Cat. 5301, ScienCell) to prevent further activation and seeded into a 6-well plate. 1 ml of collected supernatant was added to the seeded PSCs to activate them in response to tumor cell-specific secreted factors. PSCs were imaged 24 hours later using phase-contrast microscopy and then expanded in supernatant for one passage. Cells were then expanded in CAF-supplemented medium composed of a 1:1 ratio of DMEM F-12 (Cat. 31331093, Thermo Fisher Scientific): DMEM 1g/L glucose (Cat. 11885084, Thermo Fisher Scientific), 20% FBS (Cat. A5256701, Thermo Fisher Scientific), and 1% Penicillin/Streptomycin cocktail (Cat. 15070063, Thermo Fisher Scientific).

### 2D cell proliferation assay

Confluent 2D murine cells were harvested, counted, and seeded (1000 cells/well) in a 96-well plate in supplemented DMEM (4.5 g/L glucose) to a total volume of 100 µl. Proliferation was measured daily for 5 consecutive days via an MTT (3-(4,5-dimethylthiazol-2-yl)-2,5-diphenyltetrazolium bromide) colorimetric assay. Briefly, 10 µl of MTT reagent was added to each well, and the cells were incubated at standard culture conditions for 4 hours. After incubation, a solubilizing agent (50% Ethanol + 50% dimethyl-sulfoxide) was added to the wells. Solubilized formazan was detected by measuring the absorbance at 590 nm using a spectrophotometer (NanoDrop 1000, Thermo Fisher Scientific).

### RNA isolation

Cells were cultured to 80% confluency, collected, and pelleted. RNA isolation was performed using the Qiagen RNeasy mini kit (Cat. 74104, Qiagen). Briefly, the lysis buffer was prepared by adding 2-mercaptoethanol to the RLT lysis buffer (1:100). 350 µl of lysis buffer was added to confluent cells. Cells were scraped with a cell scraper or thoroughly dissociated by pipetting. Lysates were homogenized using a 21G syringe and processed immediately or stored at -80 °C until RNA isolation, as per the manufacturer’s protocols. RNA concentration was measured using a NanoDrop 1000 (Thermo Fisher Scientific).

### Reverse transcription and quantitative PCR

1μg of RNA from h/mPSCs and PSC-derived h/mCAFs was reverse transcribed to cDNA using the SensiFAST™ cDNA Synthesis Kit (Cat. BIO-65054, Meridican Biosciences). qPCR reactions were prepared using the SensiFast™ SYBR Hi-Rox Kit (Cat. BIO-92020, Meridian Biosciences) and performed on the StepOnePlus™ real-time PCR system (Thermo Fisher Scientific) according to the manufacturer’s protocols. Gene expression was quantified using the following primers: Human *18S* Forward 5’-CTCAACACGGGAAACCTCAC-3’; Human *18S* Reverse 5’-CGCTCCA CCAACTAAGAACG-3’; Human *ACTA2* Forward 5’-CTATGCCTCTGGACGCACAACT-3’; Human *ACTA2* Reverse 5’-CAGATCCAGACGCATGATGGCA-3’; Human *COL1A1* Forward 5’-TCTGCGACAACGGCAAGGTG-3’; Human *COL1A1* Reverse 5’-GACGCCGGTGGTTTCTTGGT- 3’; Murine *18S* Forward 5’-CGGACACGGACAGGATTGAC-3’; Murine *18S* Reverse 5’-TGCCA GAGTCTCGTTCGTTATC-3’; Murine *ACTA2* Forward 5’-ACTGGGACGACATGGAAAAG-3’; Murine *ACTA2* Reverse 5’-GTTCAGTGGTGCCTCTGTCA-3’; Murine *COL1A1* Forward 5’-GAGAGCGAGGCCTTCCCGGA-3’; Murine *COL1A1* Reverse 5’-GGGAGCCAGCGGGA CCTTGT-3’. Melt curve analysis was performed to ensure primer specificity and absence of primer-dimer formation. Gene expression was normalized to housekeeping genes and analyzed using the 2^−ΔΔCt method.

### Scratch Assay

Murine and PDO-derived 2D cells were harvested and counted at 80-90% confluency. A total of 100,000 cells per well were seeded in a 12-well plate using DMEM with 4.5 g/L glucose (Cat. 11965092, Thermo Fisher Scientific) or RPMI medium (Cat. 11875093, Thermo Fisher Scientific), supplemented with 10% Fetal Bovine Serum (FBS) (Cat. A5256701, Thermo Fisher Scientific) and 1% Penicillin/Streptomycin cocktail (Cat. 15070063, Thermo Fisher Scientific). Once a confluent monolayer formed, a line of cells was scraped from the center of the well to create a wound. Reference points were marked for imaging. Phase contrast microscopy was conducted at the marked areas at 0-, 4-, 8-, and 24-hours post-scratch using a Leica DM IL LED Fluo light microscope with LAS X software (version 3.5.5.19976, Leica). The acquired images were analyzed using the ImageJ Wound Healing Size Tool plugin to assess scratch closure.

### Invasion Assay

Invasion assays were performed using Matrigel Invasion Chambers with 8.0 *μ*m pore size polycarbonate membranes (Corning, Cat. #354480). Briefly, the transwell inserts were thawed at room temperature for 20 minutes and rehydrated with serum-free medium for 5 hours at 37° C prior to cell seeding. To evaluate the invasive capacity of KPC-derived cells and 2D cell lines derived from PDOs in response to fibroblast signaling, pancreatic stellate cells (mPSCs/hPSCs) or their corresponding tumor-activated PSCs (1×10^5^) or CAM tissues of similar sizes harvested at EDD9 were seeded into the lower receiver well to serve as chemo attractants. Concurrently, tumor cells (1×10^4^) cells resuspended in serum-free basal medium were seeded directly into the upper insert chamber, and the assembled plates were incubated at 37° C under sterile conditions for 24 hours.

To assess and quantify the invasive cells, crystal violet staining was performed. Following incubation, the inserts were washed with PBS, and the non-invading cells remaining on the upper surface of the membrane were gently removed with a cotton swab. The membranes were then fixed with 100% ice-cold methanol for 10 minutes at -20°C. Immediately following fixation, the inserts were incubated with a 0.5% crystal violet solution for 10 minutes at room temperature. After staining, the inserts were thoroughly washed with distilled water to remove excess dye and stored in PBS until imaging. Brightfield microscopy was performed using a Leica DM IL LED Fluo light microscope equipped with LAS X software (version 3.5.5.19976, Leica). The acquired images were analyzed using the ImageJ Cell Counter plugin to quantify the number of successfully invading cells across representative fields per membrane.

### Chick embryo chorioallantoic membrane (CAM) xenograft model preparation

CAM assay was performed as previously described with a few modifications (Ranjan et al., 2023). Briefly, fertilized specific pathogen-free (SPF) chicken eggs were obtained from ValoBioMedia, Germany, and incubated at 37 °C with 70-80% humidity for embryogenesis. On embryonic development day (EDD) 4, the eggs were turned with the rounded pole facing upwards to facilitate embryo movement, and a small window was created on the eggshell after disinfection with 70% ethanol. The window was sealed with silk tape, and the eggs were further incubated until EDD 7, when the window was enlarged for *in ovo* manipulations. Then, the window was resealed, and the eggs were incubated until EDD 9 for continued CAM development.

### CAM transplantation

On EDD 9, confluent 2D murine cells were harvested and counted. 1×10^6^ cells were resuspended in 40 µl Matrigel (Cat. 354230, Corning Life Sciences), seeded in a petri dish as Matrigel droplets, incubated at room temperature for 10 minutes, then incubated under standard cell culture conditions for 30 minutes. For PDOs, to account for varying growth rates and the time required to form organoid structures after trypsinization, single cells were seeded, depending on their growth rate, 10 days before transplantation. To transplant 2D murine cells or PDOs, the respective Matrigel droplets were scooped using a sterile spatula and dropped on the CAM. Silk tape was used to reseal the window until EDD 14.

### Harvesting of the primary tumor and chick embryo livers

On EDD 14, the primary tumor and chick embryo livers were collected. The silk tape covering the eggshell window was removed, and the CAM surrounding the tumor was cut with scissors and placed in an embedding cassette for fixation in 4% PFA overnight at room temperature. The eggshell was then cut in half, and the chick embryo was euthanized by decapitation for the collection of the liver. Chick embryo livers were transferred to cryovials, snap-frozen in liquid nitrogen, and stored at -80 °C for further processing.

### Histological Analyses

Harvested primary tumor tissue was fixed in 4% PFA overnight at room temperature. The tissue was then dehydrated and embedded in paraffin. 2.5 μm formalin-fixed paraffin-embedded (FFPE) sections were cut from each paraffin block and mounted on labeled clean adhesive microscopy slides. Hematoxylin and eosin (H&E) were used to confirm tumor architecture. Briefly, FFPE tissue sections were deparaffinized in xylene, rehydrated through a graded ethanol series, and rinsed in tap water. Nuclei were stained with hematoxylin for 30 seconds, followed by eosin staining for 20 seconds to highlight the cytoplasm. After dehydration in ethanol and clearing in xylene, the sections were mounted with Pertex mounting medium, dried overnight at room temperature, and scanned with the Aperio Versa 8 Digital Scanner for documentation using Aperio Image Scope software. Certified human pathologists at TUM Klinikum rechts der Isar reviewed all H&E sections for tumor architecture and comparisons.

### *In ovo* take-rate determination

*In ovo* take rate was determined for each PDO and the 2D murine line used in this study. To do so, the number of engrafted tumors was divided by the total number of eggs used per PDO and per 2D murine line, yielding the percentage of successfully transplanted eggs.

### Immunohistochemistry

FFPE tissue sections were deparaffinized, rehydrated, and subjected to heat-mediated antigen retrieval using a citric acid-based solution in a microwave. To minimize non-specific signals, endogenous peroxidase activity was quenched with 3% hydrogen peroxide, followed by Avidin and Biotin blocking. After blocking with 5% BSA, the sections were incubated with the following primary antibodies overnight at 4 °C: Alpha smooth muscle actin (1:500, Cat. F3777, Sigma Aldrich), Cytokeratin-19 (1:500, Cat. ab52625, Abcam), Hyaluronan Binding protein (2 μg/ml, Custom protein, R&D Systems), Ki67 (1:250, Cat. ab16667, Abcam), chicken specific Vimentin (1:250, Cat. AMF-17b, Developmental Studies Hybridoma Bank, University of Iowa). Following overnight incubation, the following biotin-conjugated secondary antibodies were added and incubated for 30 minutes at room temperature: Goat Anti-Rabbit IgG Antibody (H+L) (1:250, BA-1000, Vector Labs) or Goat Anti-Mouse IgG Antibody (H+L) (1:250, BA-9200, Vector Labs). The signal was detected using the Vectastain® elite HRP ABC reagent and developed with Vector® DAB substrate, followed by a hematoxylin counterstain. After dehydration and mounting, the slides were scanned with the Aperio Versa 8 Digital Scanner, and immunohistochemistry quantification was performed using Aperio ImageScope software with the Positive Pixel Count v9 algorithm.

### Sirius Red Staining

Sirius red staining was performed on FFPE tissue sections to quantify fibrillar collagen deposition in CAM tumors. After deparaffinization and rehydration, sections were immersed in Sirius red solution for 1 hour, covered from light. The sections were mounted and dried overnight after rinsing in absolute ethanol. The stained slides were scanned using the Aperio Versa 8 Digital Scanner. Polarized light microscopy using a DMI8 Leica Thunder microscope was used to detect collagen birefringency. The Sirius Red-positive area was quantified in Adobe Photoshop (version 22.4).

### Immunofluorescence

FFPE sections were deparaffinized, and heat-mediated antigen retrieval was performed. Next, the slides were cooled down at RT for 20-30 minutes. PDOs and 2D cells were cultured in Ibidi µ-Slide 8 and 2 Well slides, respectively (Cat. 80826 and 80286, Ibidi) until 80% confluency, fixed with 4% PFA at RT for 10-15 minutes, and stored at 4 ℃ until staining. The slides were blocked with 5% BSA in Tris-Buffered Saline with Tween 20 (TBST) for 1 hour at RT to reduce non-specific background. Overnight incubation at 4 ℃ with the following primary antibodies diluted in 5% BSA was performed: Alpha smooth muscle actin (1:500, Cat. F3777, Sigma Aldrich), Cytokeratin-19 (1:500, Cat. ab52625, Abcam), Hyaluronan Binding protein (2 μg/ml, Custom protein, R&D Systems), chicken specific Vimentin (1:100, Cat. AMF-17b, Developmental Studies Hybridoma Bank, University of Iowa) and Anti-IL-6 [1.26.4] Antibody (1:100, Cat. EUS026, Kerafast, USA). After incubation, the slides were washed with 1X TBST and incubated with the appropriate fluorescent conjugated secondary antibody: Donkey anti goat Alexa Fluor™ 488 (1:250, Cat. A-11055, Thermo Fisher Scientific), Donkey anti mouse Alexa Fluor™ 488 (1:250, Cat. A-21202, Thermo Fisher Scientific) or Donkey anti rabbit Alexa Fluor™ 594 (1:250, Cat. A-21207, Thermo Fisher Scientific). For Phalloidin staining, pre-conjugated Phalloidin-Atto 647 (1:250, Cat. 65906, Sigma Aldrich) was used. Slides were washed thrice with TBST, and counterstaining was performed with DAPI for 2 minutes at RT, protected from light. Stained sections and Ibidi slides were visualized and imaged using the Leica TCS SP8 Confocal Microscope.

### Species-specific quantification of metastatic dissemination by quantitative PCR

To assess the metastatic dissemination of PDOs and KPC murine cells in chick embryo livers, species-specific qPCR targeting *Alu* and *B1* sequences was performed. *Alu* sequences, unique to the human genome and absent in avian DNA (Zijlstra et al., 2002), were amplified to detect human DNA. In contrast, *B1* sequences, specific to the murine genome (Zhang et al., 2009), were used to identify murine DNA. Using the Nucleospin genomic DNA isolation kit (Cat. 740952.50, Macherey Nagel), DNA was isolated from chick embryo liver, and the concentration and purity were measured using a Nanodrop Spectrophotometer (Nanodrop1000, Thermo Fisher Scientific). qPCR reactions were prepared using the SensiFast™ SYBR Hi-Rox Kit (Cat. BIO-92020, Meridian Biosciences) and performed on the StepOnePlus™ real-time PCR system (Thermo Fisher Scientific) according to the manufacturer’s protocols. Chicken *GAPDH* was used as an internal control and melt curve analysis was performed to assess primer dimer formation. qPCR data were analyzed using the 2^-ΔΔCt method (Pfaffl, 2001). The following primers were used for species-specific qPCR: *Alu* Forward 5’ - ACGCCTGTAATCCCAGGACTT - 3’, *Alu* Reverse 5’ - TCGCCCAGGCTGGAGTGCA - 3’; *B1 Element* Forward 5’ -CCAGGACACCAGGGCTACAGAG - 3’, *B1 Element* Reverse 5’ - CCCGAGTGCTGGGATTAAAG- 3’; Chicken *GAPDH* Forward 5’ - GAGGAAAGGTCGCCTGGTGGATCG- 3’, Chicken *GAPDH* Reverse 5’ - GGTGAGGACAAGCAGTGAGGAACG- 3.’

### Single-cell RNA-sequencing: library preparation and sequencing

*In vitro* KPC cells were counted, diluted to an appropriate cell number in ice-cold Dulbeccós PBS (DPBS), and loaded on a 10× Chromium Next GEM Chip G to generate gel beads in emulsion (GEMs). Single-cell GEM generation, barcoding and library construction were performed using 10× Chromium Single Cell 3’ v.3.1 chemistry according to manufacturer instructions. Quality and library size of resulting cDNA and generated libraries were determined on an Agilent Bioanalyzer 2100 system using the HS DNA kit (Agilent). The library was sequenced on a NextSeq 500 (Illumina), with 26 cycles for barcodes and UMIs in read1 and 58 cycles for cDNA in read2.

### Flex Single-cell library preparation, sequencing and alignment

Flex Single-cell library preparation, sequencing, and alignment have been performed as previously described (Kose et al., 2026). Briefly, for each sample, three 50 μm FFPE tissue sections were processed according to the 10x Genomics FFPE Tissue Sections protocol (CG000632_RevD). Tissue dissociation was performed using the gentleMACS Octo Dissociator workflow. Single-cell libraries were generated following the Chromium Fixed RNA Profiling protocol (CG000527_RevF, 10x Genomics). Briefly, 500,000 cells per sample were hybridized with probes and labeled with sample barcodes at 42 °C for 20 h. Following hybridization, samples were pooled and loaded into a single well of a Chromium Chip Q for GEM generation. Sample index libraries were amplified using 10 cycles of PCR.

Libraries were sequenced on an Illumina NovaSeq X Plus using a 25B flow cell with 150 bp paired end reads. Raw sequencing data were demultiplexed using bcl2fastq (v4.2.7, Illumina). FASTQ files were processed with Cell Ranger (v9.0.1, 10x Genomics) using the multi workflow and the mouse reference transcriptome (refdata-gex-GRCm39-2024-A) together with the Chromium Mouse Transcriptome Probe Set v1.1.1. Gene expression matrices were generated using default parameters unless otherwise specified.

### Preprocessing

Cells were preprocessed following current best-practice recommendations for sc-RNA sequencing analysis (Heumos et al., 2023). Putative doublets were identified and removed using Scrublet. Count matrices were subsequently library-size normalized and log1p-transformed for downstream analyses.

### Flex-/Single cell RNA sequencing downstream analysis Dimensionality Reduction and Clustering

For dimensionality reduction of in vitro samples, the top 3,000 highly variable genes were selected, followed by principal component analysis (PCA), retaining 50 components. A k-nearest-neighbor graph was constructed (k = 15), and Uniform Manifold Approximation and Projection (UMAP) embeddings were computed for visualization. The *in vivo* samples were embedded using a drVI (Moinfar & Theis, 2024) with embedding derived from an in-house reference atlas (unpublished). Leiden clustering was performed at a resolution of 0.25. Cell type annotation was performed based on the expression of canonical marker genes identified by Leiden clustering.

### Cluster Marker Genes

Differentially expressed marker genes per Leiden cluster were identified using Scanpy’s ‘rank_genes_groups’ function with the Wilcoxon rank-sum test (Wolf et al., 2018).

### Transcriptional Heterogeneity Score

To quantify intratumoral transcriptional heterogeneity, each sample–condition combination was processed independently, such that the PCA reflects the intrinsic transcriptional variance of each population. A metacell centroid was defined as the mean PCA embedding across all cells within that group, and the heterogeneity score for each cell was computed as the Euclidean distance to its group centroid.

### PDAC Subtype Gene Signature Scoring

Published PDAC transcriptional subtype signatures, including those from Moffitt (Classical, Basal-like), Collisson (Classical, Exocrine-like, Quasi-mesenchymal), and the Bailey (Progenitor, Squamous), gene sets, were obtained from a curated gene list and converted from human to mouse gene nomenclature by capitalizing only the first letter of each gene symbol (Bailey et al., 2016), (Collisson et al., 2011; Moffitt et al., 2015). Gene set scores were computed using Scanpy’s ‘score genes’ function.

### Pseudobulk Differential Expression Analysis

To identify transcriptional differences between *in vivo* GEMM tumor cells and *in vitro* 2D cultures, pseudobulk profiles were generated for each sample by randomly partitioning cells into three pseudoreplicates (minimum 50 cells per replicate) and summing raw counts within each replicate. Pseudobulk differential expression was performed using DESeq2 (via pyDESeq2) (Love et al., 2014; Muzellec et al., 2023)

### Over-Representation Analysis

Pathway over-representation analysis (ORA) was performed on genes significantly upregulated (log₂FC > 2.0, padj < 0.05) and downregulated (log₂FC < −2.0, padj < 0.05) in GEMM tumor cells relative to 2D cultures using the Enrichr API via the gseapy Python package (Fang et al., 2023). The following gene set libraries were queried against the mouse genome: MSigDB Hallmark 2020, GO Biological Process 2023, WikiPathways 2024 (Mouse), and Reactome Pathways 2024. Pathways with an adjusted p-value < 0.05 were considered significant.

### Cell–Cell Communication

Ligand–receptor-mediated cell–cell communication was inferred using CellChat (Jin et al., 2025) with the mouse CellChatDB ligand–receptor database. Overexpressed ligands and receptors were identified using CellChat’s default method; communication probabilities were estimated using the trimean (triMean) approach; and interactions supported by fewer than 10 cells in either the sending or receiving population were filtered out. Communication probabilities were subsequently aggregated at the signaling pathway level.

### Xenium In Situ Profiling

Xenium In Situ (10X Genomics) experiments were performed using a Xenium panel comprising 377 pre-designed and 100 custom genes (Supplementary Table 2). Tissue sectioning and sample preprocessing were performed according to the manufacturer’s protocol. Briefly, 5 μm FFPE sections were mounted on Xenium slides and dried to optimize adherence. Subsequently, the samples underwent a 3-day protocol comprising multiple steps, including deparaffinization, decrosslinking, probe hybridization, ligation, amplification, and cell segmentation staining.

Preprocessed samples were analyzed using the Xenium Analyzer with instrument software version 4.0.1.4 and analysis version 4.0.1.0. Subsequently, H&E staining of the tissue sections was performed according to the manufacturer’s protocol for post-Xenium stainings. Following H&E staining, slides were scanned using an Aperio Imager AT2 (Leica, Germany) at 40× magnification (pixel width, 0.253 µm).

### H&E Destaining

Prior to ALU-ISH staining, the H&E stain was removed from the tissue sections. Coverslips were detached by immersion in xylene overnight. Sections were then rehydrated through a descending alcohol series (isopropanol × 2, 96% ethanol × 2, 70% ethanol × 2; 2–3 min each step). Hematoxylin was extracted by incubation in 0.75% HCl in ethanol (HCl-Alkohol, Carl Roth) until no residual staining was visible to the naked eye. Sections were washed three times in distilled water.

### ALU In Situ Hybridization (ALU-ISH)

ALU-ISH was performed on the BOND RX system (Leica, Germany), an automated staining platform, using the Bond ALU-ISH protocol (all reagents from Leica). Slides were deparaffinized (3 min) followed by alcohol treatment to wash buffer (20 min). Endogenous peroxidase activity was blocked using Peroxide Block (5 min). Enzymatic pretreatment was performed with Enzyme 1 (Leica; 15 min), followed by denaturation (10 min). Hybridization with an ALU probe was carried out for 2 hours. Stringency washing was performed at 37 °C for 10 min using Stringency Wash Solution (Leica). The samples were incubated with an anti-biotin antibody (Leica; 30 min), Post Primary reagent (Bond Polymer Refine Kit, Leica; 8 min), and Polymer (Bond Polymer Refine Kit, Leica; 8 min), followed by chromogenic detection with mixed DAB Refine (10 min). Counterstaining was performed with hematoxylin (5 min). Wash buffer steps were applied between each incubation step. Stained slides were scanned using an Aperio Imager AT2 (Leica, Germany) at 40X magnification (pixel width of 0.253 µm).

### Segmentation and Quality Control

To improve cell segmentation quality, we applied Proseg (Jones et al., 2025) (v3.1.0) using the parameters—xenium and --enforce-connectivity, with min_qv = 20, --diffusion-probability = 0.0, and --max-transcript-nucleus-distance = 20. Transcripts matching the following patterns were excluded: *Deprecated*, *NegControl*, *Unassigned*, *Intergenic*, *BLANK*, and antisense transcripts. The raw transcripts.csv.gz files served as input. Image registration was performed using the InSituPy package (Jones et al., 2025). Following segmentation, standard single-cell quality control was performed. Cells with fewer than 15 detected gene (n_genes_by_counts < 15) or fewer than 30 total counts (total_counts < 30) were excluded from further analysis. Data normalization was performed using a target sum of 500 counts per cell to account for the characteristics of Xenium *In Situ* (XIS) sequencing, followed by logarithmic transformation.

### Batch Correction and Cell Type Annotation

To account for batch effects, we applied Harmony (Korsunsky et al., 2019) integration using 50 principal components, with sample type and sample ID specified as batch covariates. Primary cell type annotation was performed using a data-driven approach combined with canonical marker genes. Subsequently, cell type annotations and the batch-corrected latent representation were refined using resolVI (Ergen & Yosef, 2025). Tumor subtype annotation was based on Moffitt signatures (Moffitt et al., 2015) and scored using Scanpy’s score_genes function (Wolf et al., 2018). Scores were z-score normalized, and differential scores between subtype signatures were computed. Cells within the top 33% of the differential score between classical and basal signatures were assigned to their respective subtypes, while the remaining cells were classified as intermediate. For the Bailey (Bailey et al., 2016) and Puleo (Puleo et al., 2018) subtype classifications, assignments were based on differences in signature scores, with a threshold of 0.5. CAFs were further characterized by scoring iCAF and myCAF signatures using the *ulm* method implemented in decoupler (Badia et al., 2022). Cells were assigned to a subtype based on the highest score, considering both score magnitude and statistical significance (p-value).

### Distance-Based Heterogeneity

To quantify tumor cell heterogeneity between patient samples and matched PDO-CAM models, we computed an average tumor cell for each sample in the shared 50-dimensional Harmony embedding. Euclidean distances were then calculated between each tumor cell and the corresponding sample-specific centroid. To assess the effect size of differences between groups, we applied Cliff’s delta. Proliferative tumor cells were defined based on *MKI67* expression, with cells above the 70th percentile of *MKI67*-expressing cells classified as proliferative.

### Data Availability

Raw data from scRNA seq and spatial transcriptomics have been submitted to GEO: SUB16275073. Due to the potential re-identifiability and to prevent personal data and privacy issues, we are unable to make raw genomic (DNA) sequencing data available (Shabani & Marelli, 2019).

### Software

All bioinformatic analyses were performed in Python using InSituPy, Scanpy, AnnData, harmony-pytorch, and decoupler, along with standard scientific libraries (NumPy, Pandas, SciPy, Seaborn, Matplotlib, and cliffs-delta). Statistical analyses were performed using GraphPad Prism (Version 11.0.1).

## Results

### Single-cell transcriptomic analysis reveals loss of phenotypic heterogeneity and convergence of KPC tumor cell states in 2D culture

Single-cell transcriptomic profiling was performed on three KPC-derived pancreatic cancer cell lines, 9366, 12548, and R405, together with their matched parental KPC tumors from which the cell lines were established (Supplementary Figure 1A). Comparative analysis of malignant cells derived from 2D cultures and corresponding primary tumors revealed marked differences in their transcriptional landscapes. Malignant cells from primary tumors exhibited greater transcriptional heterogeneity than cultured cells, suggesting that the in *vivo* environment supports a broader spectrum of cellular states and gene expression programs that are not fully maintained under standard 2D culture conditions (Figure 1A–C). Quantitative assessment of distances to cluster centroids revealed significantly greater transcriptional dispersion in KPC tumors compared with matched KPC cell lines (Figure 1D). Density and cumulative distribution analyses confirmed a global shift toward larger centroid distances in tumors, indicating increased transcriptional heterogeneity among malignant cells *in vivo (*Supplementary Figure 1B).

**Figure 1.**
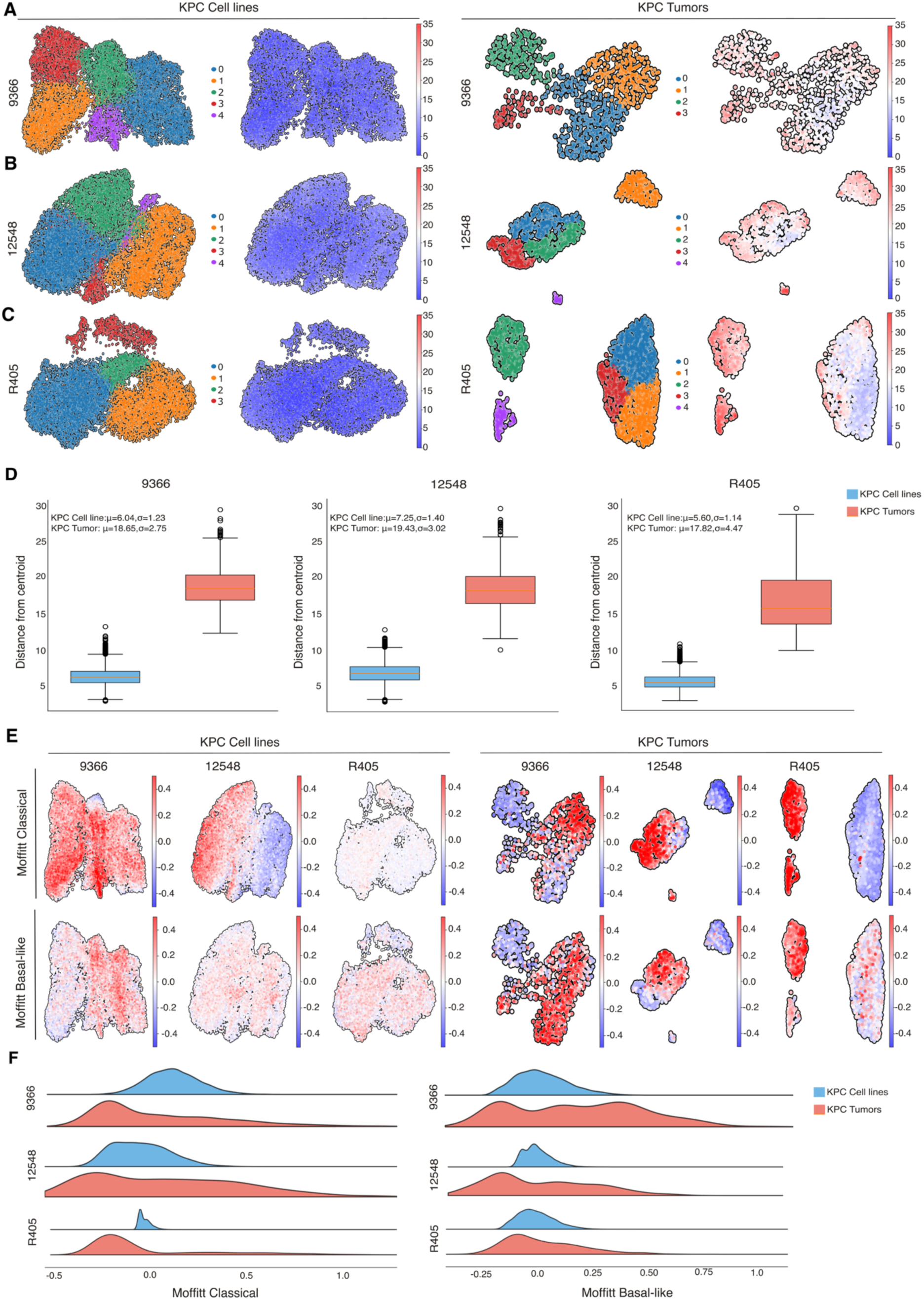
Single-cell transcriptomic profiling reveals cellular heterogeneity and PDAC subtype distributions in KPC-derived cell lines and matched tumors., (A–C) UMAP visualizations of scRNA-seq data from KPC cell lines (left) and their corresponding parental KPC tumors (right): (A) 9366, (B) 12548, and (C) R405. Cells are annotated by unbiased Leiden clusters (left) and by distance from the cluster centroid (right). (D) Box plots comparing the distribution of cell-to-centroid distances between each cell line and its matched parental tumor. (E) UMAP projections annotated according to PDAC subtype scores derived from the Moffitt classification framework, illustrating subtype-specific transcriptional programs in KPC cell lines (left) and KPC tumors (right). (F) Ridgeline plots showing the distribution of Moffitt PDAC subtypes in KPC cell lines and KPC tumors.

Transcriptional programs retained *in vitro* converged toward relatively homogeneous cellular states marked by the expression of proliferation-associated genes (Supplementary Figure 1C). By contrast, tumor-derived cells displayed greater transcriptional heterogeneity and expression of genes linked to mesenchymal features, ECM interactions, stress adaptation, and cellular plasticity (Supplementary Figure 1D). These findings indicate that tumor cells *in vivo* occupy a broader spectrum of transcriptional states than their corresponding cell lines and suggest that some of this diversity may depend on signals present within the native TME.

Comparative analysis across KPC cell lines (Supplementary Figure 1E) and matched KPC tumors (Supplementary Figure 1F) further demonstrated variable degrees of phenotypic stability and plasticity. Although tumor-specific lineage-associated transcriptional programs were partially retained following adaptation to culture conditions, 2D propagation substantially constrained transcriptional diversity and promoted convergence toward restricted epithelial and proliferative states.

Projection of previously published PDAC subtype signatures (Bailey et al., 2016; Moffitt et al., 2015; Puleo et al., 2018) onto the KPC models further revealed pronounced subtype convergence *in vitro*. Specifically, the 9366, 12548, and R405 cell lines predominantly exhibited classical, mixed, and basal-like phenotypes, respectively (Figure 1E-F; Supplementary Figure 2A–D). In contrast, matched KPC tumors displayed marked intratumoral subtype heterogeneity, with coexistence of multiple PDAC transcriptional subtypes within individual tumors (Figure 1E-F; Supplementary Figure 2A–D).

Collectively, these findings demonstrate that adaptation to 2D culture conditions reduces malignant cell-state diversity and drives transcriptional convergence toward restricted subtype-associated phenotypes, whereas primary tumors maintain heterogeneous cellular programs through dynamic interactions with the TME.

### KPC tumor cells re-establish subtype-specific tumor architecture and stromal organization within an *in ovo* niche

Guided by our single-cell transcriptomic analyses demonstrating that conventional 2D culture promotes transcriptional convergence and loss of malignant-cell heterogeneity, we next investigated whether exposure to a naïve *in ovo* microenvironment could restore tumor architecture and phenotypic diversity. To address this question, we established a KPC-on-CAM platform by engrafting KPC cell lines onto the CAM using the experimental workflow illustrated in Figure 2A. Transplantation of 1 × 10^6^ cells resulted in robust and reproducible tumor engraftment within five days (Supplementary Figure 3A).

**Figure 2.**
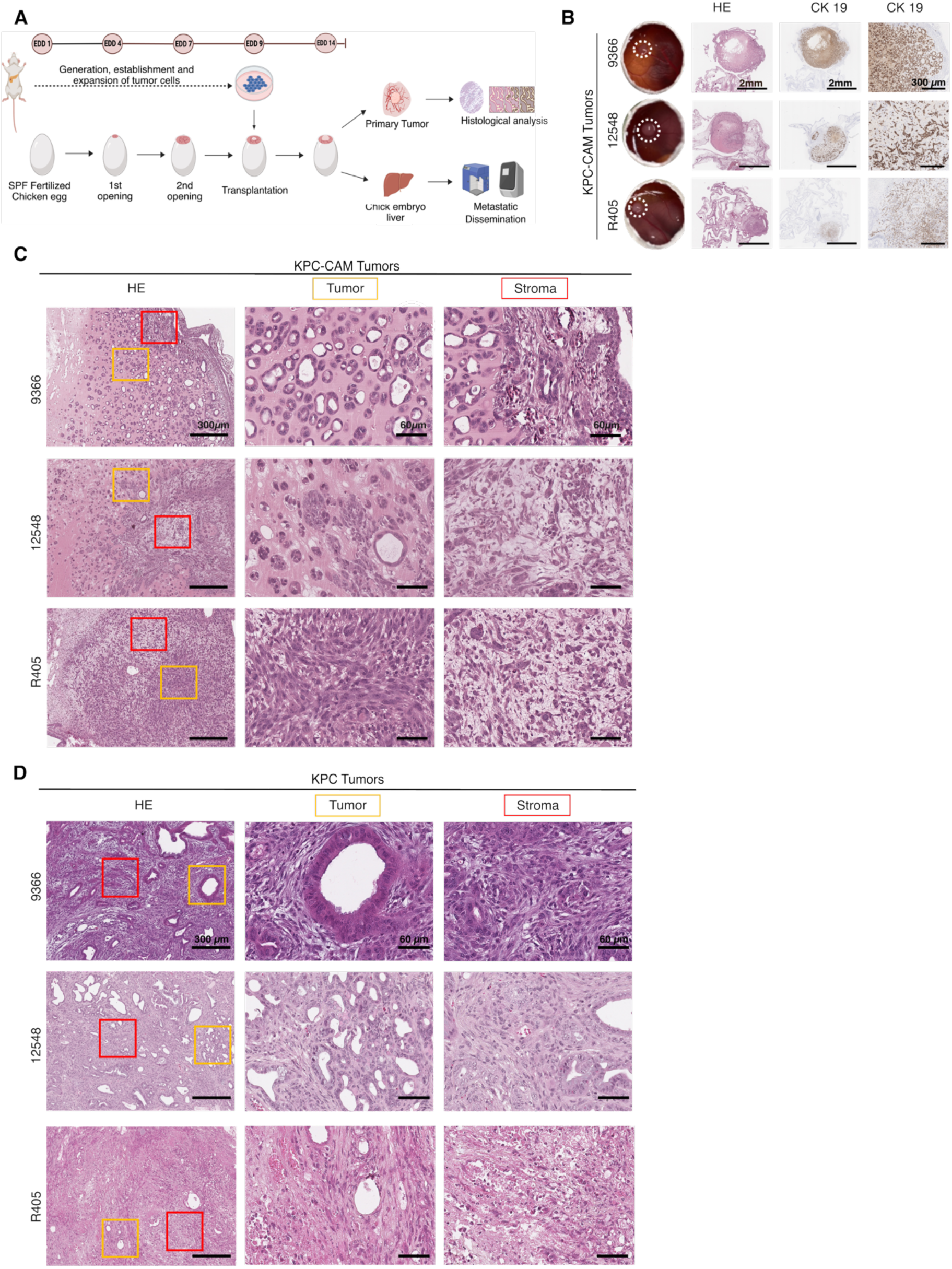
Establishment and histomorphological analysis of KPC-on-CAM tumors (A) Schematic representation of the KPC-on-CAM model establishment. Fertilized SPF eggs were incubated, and on embryonic development day (EDD) 4, a small window was opened to separate the CAM from the eggshell membrane. The window was expanded on EDD 7. On EDD 9, 1× 10⁶ murine KPC cells were resuspended in Matrigel and transplanted onto the CAM. Primary tumors and chick embryo liver were harvested on EDD 14 for downstream analyses. (B) Representative images of primary tumors formed on the CAM (left) and an overview of H&E and CK19 staining of KPC-CAM tumors at 2X magnification. Scale bar = 2 mm. CK19 staining of KPC-CAM tumors at 10X magnification. Scale bar = 300 μm. (C) H&E of KPC-CAM tumors at 10X magnification. Scale bar = 300 μm. Tumor and stroma are marked in red and yellow squares, respectively. Scale bar = 60 μm. (D) H&E staining of endogenous mouse tumors at 10X magnification. Scale bar = 300 μm. Tumor and stroma are marked in red and yellow squares, respectively. Scale bar = 60 μm.

Histological characterization of KPC-CAM tumors by H&E staining, together with Cytokeratin 19 (CK19) immunohistochemistry, confirmed tumor identity and enabled clear discrimination between the malignant epithelial and stromal compartments (Figure 2B). 9366-CAM tumors predominantly exhibited moderately differentiated ductal morphology. In contrast, 12548-CAM tumors displayed mixed regions of moderately and poorly differentiated cells, associated with variable stromal responses. R405-CAM tumors demonstrated predominantly poorly differentiated growth accompanied by extensive stromal infiltration and a prominent myxoid stromal architecture (Figure 2C). These histomorphological features were consistent with the phenotypic characteristics of the corresponding parental cell lines and were further supported by F-actin staining patterns (Supplementary Figure 3B). Comparison with matched endogenous mouse tumors demonstrated substantial preservation of histopathological characteristics across models (Figure 2C-D). Endogenous 9366 tumors exhibited moderately differentiated ductal growth with prominent desmoplastic stroma, whereas 12548 tumors showed mixed differentiation states accompanied by stromal remodeling. Similarly, endogenous R405 tumors displayed poorly differentiated morphology, extensive stromal infiltration, and additional areas of necrosis *in vivo* (Figure 2D).

Collectively, these findings demonstrate that KPC-on-CAM xenografts faithfully recapitulate key histomorphological features of endogenous PDAC tumors while preserving subtype-associated phenotypic characteristics observed *in vitro.* This observation was consistent upon engrafting the morphologically characterized KC murine cell line onto the CAM (Supplementary Figure 3C–E). Moreover, the KPC-on-CAM model retained tumor-grade-associated architectural features corresponding to the matched endogenous PDAC tumors (Table 1), supporting the ability of the *in ovo* microenvironment to reestablish complex tumor organization and phenotypic diversity.

**Table 1.**
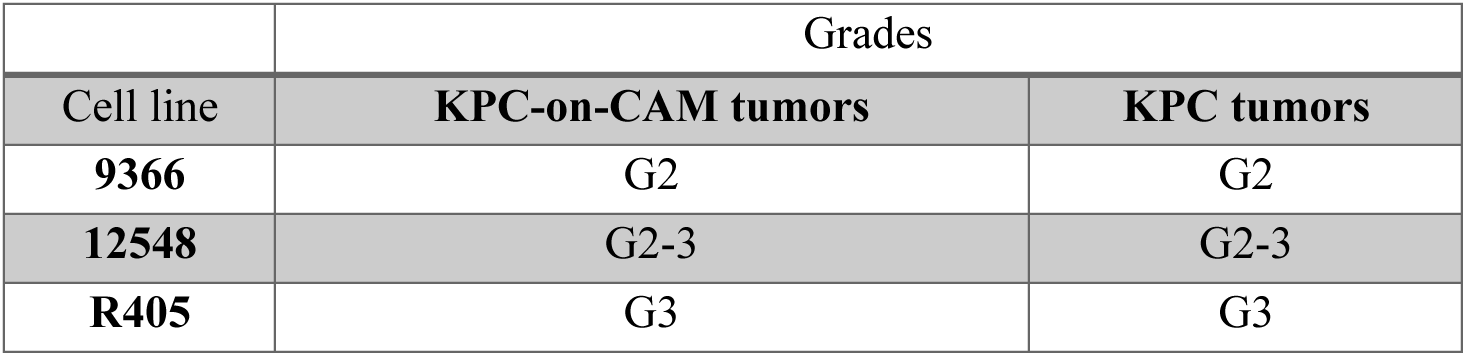
Tumor grades of KPC-on-CAM and corresponding endogenous mouse tumors.

### Active stromal remodeling by KPC tumor cells recapitulates key features of the pancreatic tumor microenvironment *in ovo*

To investigate how these baseline tumor profiles dictate the composition of the surrounding stroma, we leveraged our endogenous single-cell datasets from KPC tumors (9366, 12548, and R405) to evaluate host stromal composition (Figure 3A-B). Cell-cell communication and network analysis via CellChat revealed that the malignant epithelial compartment acts as a central communication hub, displaying the highest number of overall signaling interactions across all cell types in the TME (Figure 3C). Crucially, evaluation of interaction strength demonstrated that the most robust, high-intensity crosstalk occurred directly between malignant cells and CAF, namely the myofibroblastic (myCAF) and inflammatory CAF (iCAF) subpopulations (Figure 3D).

**Figure 3.**
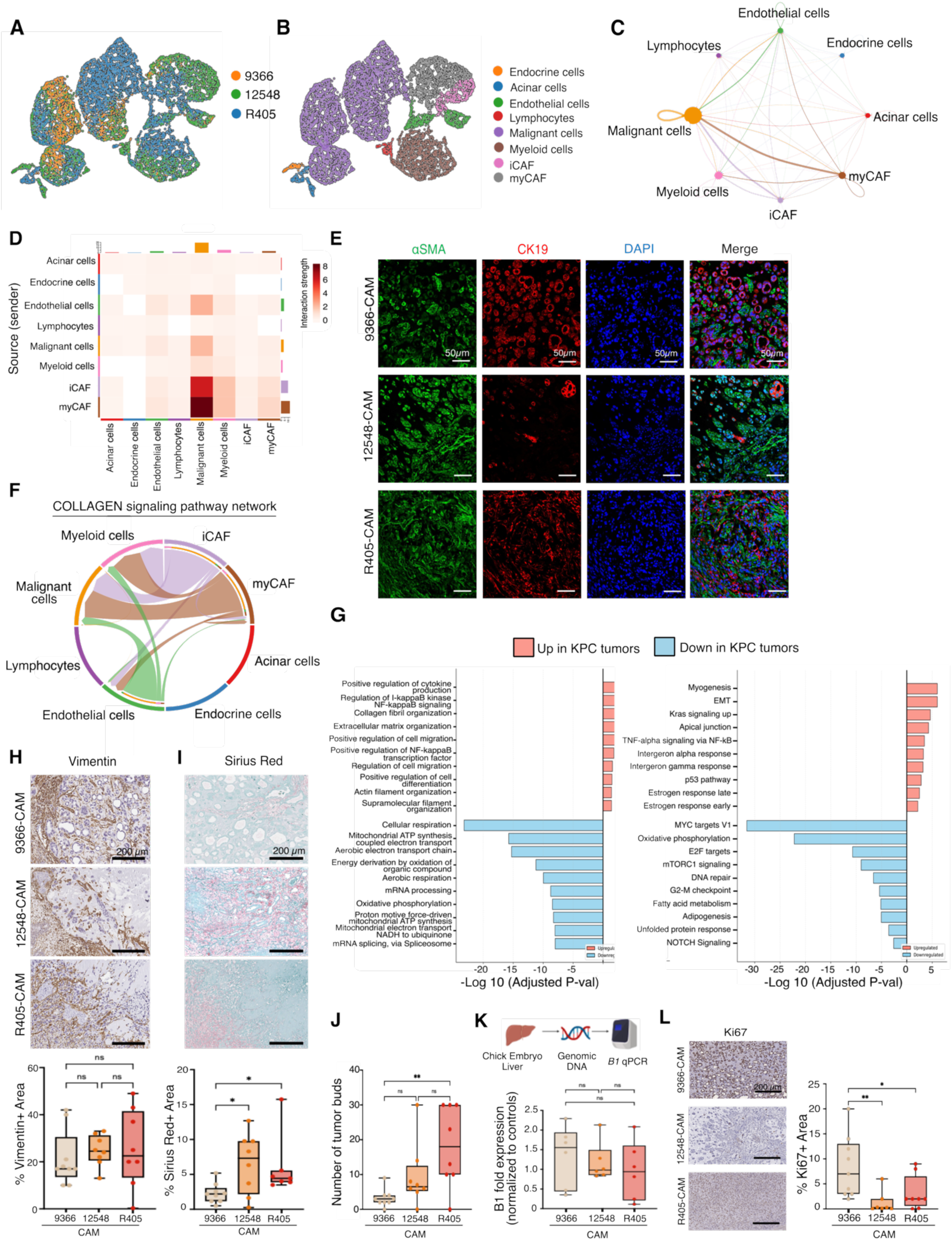
Active stromal remodeling by KPC cells recreates the pancreatic TME *in ovo*. (A) UMAP visualization of KPC tumors (9366, 12548, R405). (B) Cell type annotation showing specific populations, including malignant cells, myeloid cells, inflammatory CAFs (iCAFs), and myofibroblastic CAFs (myCAFs), within endogenous mouse tumors. Network cell-cell communication plot (CellChat) mapping (C) interaction frequencies and (D) interaction strength. (E) Representative immunofluorescence images at 40X magnification showing activated αSMA-positive stroma (green), malignant epithelial structures marked by CK19 (red), and nuclei counterstained with DAPI (blue) across KPC-CAM tumors. Scale bar = 50 μm. (F) Chord diagram showing collagen associated pathway interactions between tumor and TME compartments. (G) Gene Ontology (GO) biological pathways (left) and MSigDB Hallmark (right) enrichment analysis between the malignant epithelial compartments of KPC tumor and KPC cell lines. Data presented as pathways upregulated (red) or downregulated (blue) in KPC tumor. (H) Representative immunohistochemical staining (at 20X magnification) and quantification avian fibroblasts via a chicken-specific Vimentin antibody. Scale bar = 200 μm. (I) Representative images (at 20X magnification) and matching quantitative analysis of Sirius Red staining, demonstrating variations in host collagen matrix deposition (*p=0.0159; 0.0137). Scale bar = 200 μm. (J) Quantitative analysis of malignant tumor budding identified at the active peripheral edges of the KPC-CAM tumors (**P=0.0079). (K) *B1* qPCR analysis of chick embryo livers. (L) Representative immunohistochemistry images (at 20X magnification) and quantitative evaluation of Ki67 expression across KPC-CAM tumors. Scale bar = 200 μm. For all *in ovo* CAM experiments (H–L), sample sizes are *N*=9 for 9366 KPC-CAM, *N*=8 for 12548 KPC-CAM, and *N*=9 for R405 KPC-CAM. Statistical significance was determined by the Kruskal-Wallis test (**p=0.0017; *p=0.324). Data are presented as median with Min to Max range; ns = non-significant.

To confirm the presence of CAFs *in ovo*, we performed immunofluorescence for alpha-smooth muscle actin (α-SMA) and CK19 on KPC-CAM tumors. We observed specialized spatial patterns where highly activated α-SMA-positive stromal cells tightly encapsulated the malignant ductal structures (Figure 3E). To identify the underlying ligand-receptor systems that dictate this crosstalk, we used a rankNET hierarchical plot to rank signaling pathway information flow. This revealed multiple ECM components of interest, with the Collagen pathway displaying the highest overall communication hierarchy, followed closely by fibronectin (*FN1*), laminin, and thrombospondin (*THBS*) networks (Figure 3F, Supplementary Figure 4A).

A major challenge in utilizing cell lines for such studies is the rapid phenotypic and molecular drift induced by culture on plastic. To define this transcriptomic divergence, we performed differential pathway analysis directly comparing the malignant epithelial compartment of endogenous GEMM tumors against their 2D-cultured counterparts. This analysis revealed a profound transcriptomic shift: 2D-isolated cells down-regulated crucial pathways involved in cellular respiration, oxidative phosphorylation, and tissue architecture, while up-regulating stress- and cell-cycle-associated programs (Figure 3G). Remarkably, when these 2D cells are engrafted *in ovo*, the 3D architecture of the CAM environment acts as a corrective niche, allowing them to self-organize and recapitulate the complex glandular tumor architecture and desmoplastic landscape observed in the original GEMM tumors (Figure 2C).

We next hypothesized that these KPC lines actively recruit and reprogram naïve host cells to shape this peri-tumoral architecture. To evaluate this mechanism, we treated naïve mouse pancreatic stellate cells (mPSCs) *in vitro* with conditioned media (CM) collected from each KPC line. Upon exposure to KPC-CM, mPSCs underwent a definitive morphological transition, shifting from an inactivated morphology into a highly elongated, spindle-shaped myofibroblastic phenotype characterized by organized α-SMA expression (Supplementary Figure 4B). qPCR of these activated cells confirmed significant increases in transcript levels of canonical fibroblastic markers *Acta2* and *Col1a1* (Supplementary Figure 4C–D). Notably, lines R405 and 12548 induced the most robust transcriptional up-regulation, matching the elevated expression levels of myCAF and iCAF marker sets verified in our tracking matrices as seen in the CAFs from scRNA-seq of the corresponding GEMM (Supplementary Figure 4E).

To assess whether the tumor cell–induced fibroblast activation observed *in vitro* is recapitulated at the tissue level, we evaluated the stroma of the corresponding *in ovo* KPC tumors. Immunohistochemical staining for chicken-specific vimentin (AMF-17b) confirmed a dense influx of host avian fibroblasts enveloping the tumor cells across all lines (Figure 3H). Sirius Red staining further revealed extensive host collagen deposition across all lines (Figure 3I). Concordant with our *in vitro* mPSC activation data, lines 12548 and R405 showed significantly higher percentages of Sirius Red-positive area than line 9366, demonstrating that the intrinsic cell state of the tumor cells influences structural matrix deposition *in ovo*. This fibrous matrix remodeling was further validated by a significant increase in Hyaluronan-Binding Protein (HABP) positive areas in lines 12548 and R405 compared to 9366 (Supplementary Figure 4F). Indeed, multiplex immunofluorescence for chicken-specific Vimentin, Interleukin-6 (IL-6), and HABP validated the localized arrangement of activated and cytokine-expressing host-CAF networks directly integrated within dense hyaluronic acid matrices at the tumor-avian tissue interfaces (Supplementary Figure 4G).

Having established differences in stromal activation and ECM remodeling, we next asked whether these distinct microenvironmental states translated into functional differences in tumor cell invasion and metastatic dissemination. *In vitro* scratch-wound assays showed uniform baseline migration profiles across the lines on standard plastic (Supplementary Figure 4H-I). However, invasion assays demonstrated that the presence of mPSC derived CAF (+CAF) or CAM tissue fragments acted as powerful chemotactic triggers, driving a significant and comparable increase in the invasive potential of KPC cells (Supplementary Figure 4J).

Histopathological evaluation of the primary *in ovo* tumors confirmed that line R405, the highly mesenchymal, matrix-dense line, produced a significantly higher number of tumor buds at the active tumor edge compared to lines 9366 and 12548 (Figure 3J). Systemic dissemination to host embryo livers was measured by qPCR for the *B1* transposon sequence, a DNA element found primarily in the genomic DNA of murine but not avian cells, which confirmed early target organ dissemination across all groups (Figure 3K). Crucially, to determine whether this aggressive dissemination pattern was biased by accelerated cell proliferation, we quantified primary tumor Ki-67 expression (Figure 3L). R405-CAM showed a paradoxically lower Ki-67+ proliferative area than line 9366-CAM. This demonstrates that early metastatic dissemination and tumor budding in these models are primary functional outcomes of active, EMT-driven stromal invasion rather than an artifact of rapid tumor cell proliferation.

Overall, these parallels with established murine datasets suggest that the *in ovo* model serves as a reliable surrogate platform, capturing key features of tumor-stroma architecture and functional dynamics observed in traditional rodent models and represents a viable, complementary approach for modeling complex pancreatic tumor biology.

### PDO-on-CAM xenografts reconstruct patient-specific architectural programs of PDAC

The ability of KPC cells to self-organize into complex tumors and actively remodel the stromal niche *in ovo* suggested that the CAM platform may provide a permissive environment for pancreatic tumor reconstruction. We therefore investigated whether this approach could be translated to patient-derived models by establishing a hybrid PDO-on-CAM platform. PDOs from EUS-FNA/B specimens and surgical resections were molecularly characterized (Supplementary Table 1, Supplementary Figure 5A–B). and optimized for CAM transplantation (Figure 4A, Supplementary Figure 5C–D). A seeding density of 100 × 10^3^ cells achieved reproducible engraftment rates of ∼80% within five days across PDO lines (Supplementary Figure 5E). Engrafted tumors and chick embryo livers were subsequently analyzed for histopathology and metastatic dissemination, establishing the PDO-on-CAM system as a robust translational model.

**Figure 4.**
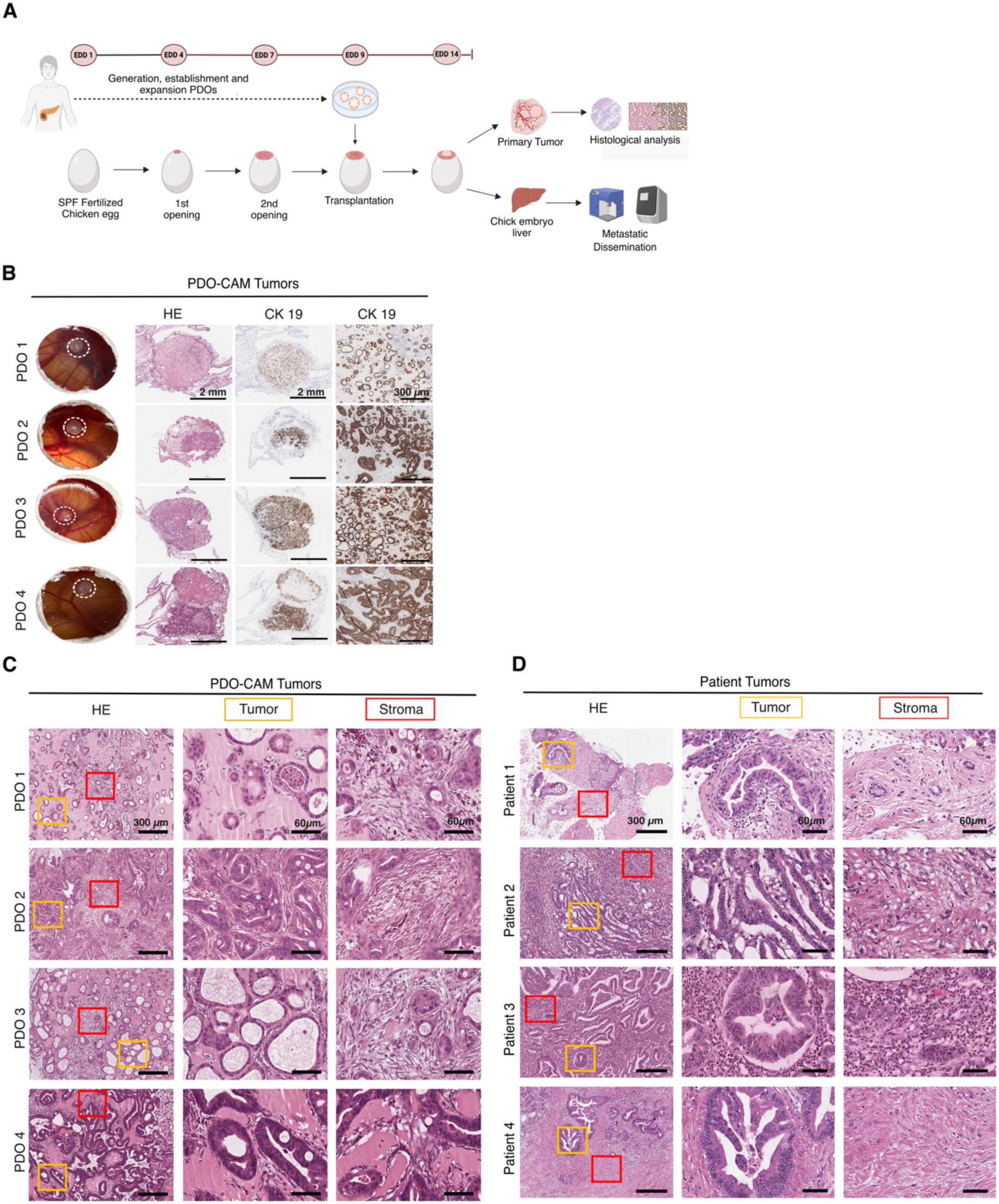
Establishment and histomorphological analysis of PDO-on-CAM tumors (A) Schematic representation of the PDO-on-CAM model establishment. On EDD 9, Matrigel domes containing 1 × 10^5^ PDO cells (seeded 10 days prior to account for varying proliferation rates) were transplanted onto the CAM. Primary tumors and chick embryo liver was harvested on EDD 14 for downstream analyses. (B) Representative images of primary tumors formed on the CAM and overview of H&E and CK19 staining of PDO-on-CAM tumors at 2X magnification. Scale bar = 2 mm. CK19 staining of PDO-on-CAM tumors at 10X magnification. Scale bar = 300 μm. (C) H&E staining of PDO- CAM tumors at 10X magnification. Scale bar = 300 μm. Tumor and stroma are marked in red and yellow squares, respectively. Tumor and stroma compartments at 40X magnification. Scale bar = 60 μm. (D) H&E staining of primary patient tumors at 10X magnification. Scale bar = 300 μm. Tumor and stroma are marked in red and yellow squares, respectively. Tumor and stroma compartments at 40X magnification. Scale bar = 60 μm.

Histological characterization of PDO-on-CAM tumors by H&E staining and CK19 immunohistochemistry confirmed ductal epithelial tumor identity and enabled clear differentiation from the surrounding stromal compartment (Figure 4B). Subsequent comprehensive histomorphological analyses revealed distinct patient-specific architectural phenotypes (Figure 4C). PDO-CAM 1 tumors displayed multifocal ductal neoplastic growth with flat-to-cuboidal tumor cell morphology, accompanied by sparse stromal infiltration, a loose myxoid stroma, and minimal immune cell presence. PDO-CAM 2 tumors exhibited ductal-to-cribriform and papillary-to-micropapillary growth patterns composed of columnar tumor cells, together with moderate ECM deposition and increased stromal activation. PDO-CAM 3 tumors predominantly showed ductal architectures with flat-to-cuboidal tumor cells embedded within loose myxoid stroma and limited inflammatory infiltration. In contrast, PDO-CAM 4 tumors demonstrated tubular-to-cribriform growth patterns, with multilayered epithelial organization and moderate stromal infiltration within a minimally reactive myxoid stroma.

To determine the extent to which PDO-on-CAM tumors recapitulate native tumor histology, corresponding primary patient tumors were comparatively analyzed (Figure 4D). Patient tumor 1 exhibited heterogeneous ductal and columnar growth patterns, moderate ECM deposition, and limited immune infiltration. Patient tumor 2 displayed highly cellular ductal and tubular architectures with multilayered tumor organization, sparse stromal content, and extensive necrotic and inflammatory regions. Patient tumor 3 demonstrated papillary, cuboidal, and columnar growth patterns associated with pronounced stromal infiltration and inflammatory cell accumulation. Patient tumor 4 exhibited large duct-like structures with papillary intraluminal projections, moderate-to-high stromal infiltration, and a loose, desmoplastic, myxoid stroma containing sparse immune infiltrates.

Overall, PDO-on-CAM tumors preserved key histomorphological characteristics of their corresponding patient tumors, including architectural organization, stromal composition, and tumor grade (Table 2). These findings demonstrate that the PDO-on-CAM platform faithfully recapitulates patient-specific pancreatic tumor morphology and supports its application as a translational model for individualized tumor biology.

**Table 2.**
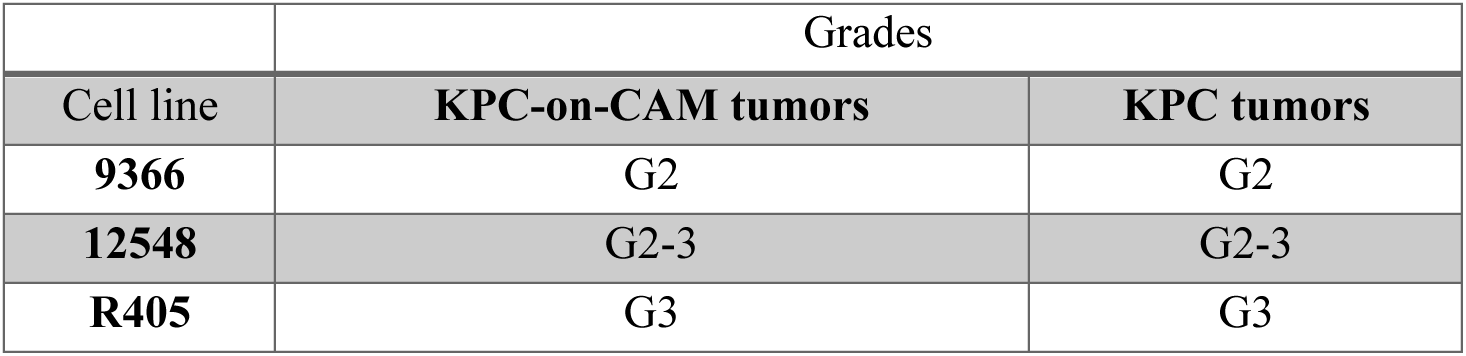
Tumor grades of PDO-on-CAM and the corresponding patient tumors.

### Spatial transcriptomic analysis reveals preservation of tumor cell heterogeneity and architectural programs in PDO-on-CAM tumors

Spatial transcriptomic profiling was conducted on patient tumors and their matched PDO-CAM tumors to assess the extent to which the PDO-CAM model preserves the cellular and molecular features of the original tumors. Patient 1 was excluded from this analysis due to limited tissue availability. Cell type annotation revealed a complex cellular landscape in patient tumors, comprising acinar cells, B/plasma cells, ductal cells, endothelial cells, macrophages, T cells, CAFs, tumor cells. In contrast, PDO-CAM tumors were predominantly composed of tumor cells. Cells that were not identifiable were labeled as mixed cells (Figure 5A-C). This apparent reduction in cellular complexity is likely attributable not only to biological differences, such as an immature avian immune system at the time of xenografting, but also to technical limitations of spatial transcriptomic analysis. Specifically, the Xenium assay employed a human-specific gene panel, which may not efficiently detect or classify host-derived chick stromal and immune cells present in the CAM microenvironment. Consequently, certain non-human cellular populations may be underrepresented or absent in the PDO-CAM datasets, potentially contributing to the observed decrease in cellular diversity.

**Figure 5.**
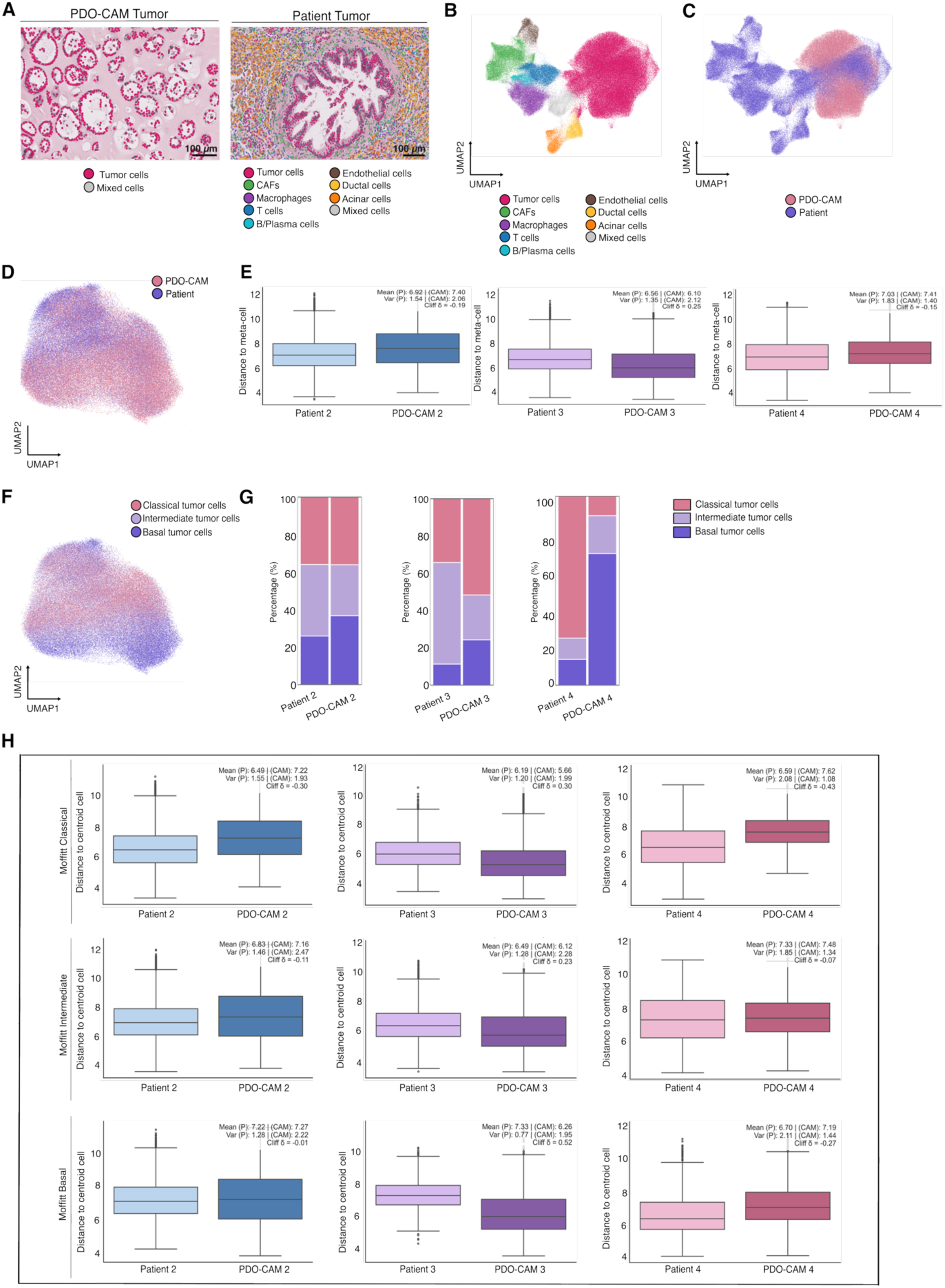
Spatial transcriptomic analysis of tumor cell states in patient tumors and PDO-CAM tumors. (A) Representative H&E tissue sections from the PDO-CAM and patient tumors used for Xenium spatial transcriptomic profiling. Cells were annotated based on transcriptomic signatures and overlaid onto the corresponding histological images. (B) UMAP visualization of all annotated cells from both patient and PDO-CAM tumors according to cell-type identity. (C) UMAP of all annotated cells by sample origin. (D) UMAP embedding of tumor cells only, extracted from both patient and PDO-CAM tumors. (E) Distribution of the distance of individual tumor cells to the corresponding metacell centroid across samples. (F) UMAP of tumor cells classified according to Moffitt molecular subtypes (Classical, Intermediate, and Basal) across both patient and PDO-CAM tumors. (G) Cell composition analysis showing the relative abundance of Moffitt tumor subtypes within each sample. (H) Box plots showing the distribution of distances from tumor cells to their subtype-specific centroid for each Moffitt subtype.

To assess the extent to which tumor-intrinsic cellular heterogeneity was retained after engraftment, we performed a focused analysis of the tumor cell compartment (Figure 5D). Distance-to-meta cell analysis revealed high concordance between tumor cell populations from patient tumors and matched PDO- CAM tumors, demonstrating that the model faithfully preserves the transcriptional landscape and cellular heterogeneity of the original tumors (Figure 5E).

To determine whether PDO-CAM tumors preserved the cellular proliferative states present in the corresponding patient tumors, we first assessed *MKI67* expression within the tumor cell compartment, which revealed the presence of highly proliferative and non-proliferative compartments (Supplementary Figure 6A). Distance-to-meta cell distributions were comparable between proliferative and non-proliferative tumor cell populations, demonstrating that the retained intra-tumoral heterogeneity was not driven solely by cell-cycle–associated transcriptional programs (Supplementary Figure 6B).

We next investigated whether tumor subtype heterogeneity was similarly conserved by scoring tumor cells using previously established PDAC subtype gene signatures (Bailey et al., 2016; Moffitt et al., 2015; Puleo et al., 2018). (Figure 5F-G, Supplementary Figure 6C-D). Spatial transcriptomic analysis revealed a comparable distribution of subtype-associated transcriptional programs between patient tumors and their corresponding PDO-CAM tumors, indicating that the molecular subtype architecture of the original tumors is largely maintained (Figure 5H and Supplementary Figure 6E-F). Together, these findings demonstrate that PDO-CAM tumors preserve both intratumoral transcriptional heterogeneity and subtype diversity, supporting their utility as a patient-representative model for studying PDAC biology.

### Spatial transcriptomics demonstrates preservation of CAF heterogeneity between patient tumors and matched PDO-CAM xenografts

Having established the spatial landscape of the malignant epithelial cells, we next turned our attention to the surrounding stromal compartment within the same Xenium dataset (Patient 2, 3, and 4). Unsupervised clustering of the stromal cells identified distinct sub-populations of CAFs (Figure 6A). Detailed annotation of these clusters revealed three distinct CAF phenotypes: CAFs, iCAFs, myCAFs (Figure 6B). Due to the focused gene panel of the Xenium V1 platform, cells expressing baseline markers of both activation states without clear polarization were classified broadly as unaligned CAFs. Intriguingly, when correlating these stromal phenotypes back to the malignant epithelial patterns described previously, we observed a distinct tissue-level polarization. Specifically, Patient tumour 4, which harbored the highest density of classical-like tumor cells corresponded to the highest proportional abundance of iCAFs within its local stroma (Figure 6C). To confirm the molecular identity of these sub-populations, we mapped the expression of canonical CAF markers across the patient UMAP space. The myCAF cluster was characterized by strong enrichment of *ACTA2*, *COL5A1*, *POSTN*, *MYH11*, and *THBS2*, whereas the iCAF cluster was distinctly marked by complement component *C3* expression (Figure 6D; additional myCAF and iCAF markers are presented in Supplementary Figure 7A-B).

**Figure 6.**
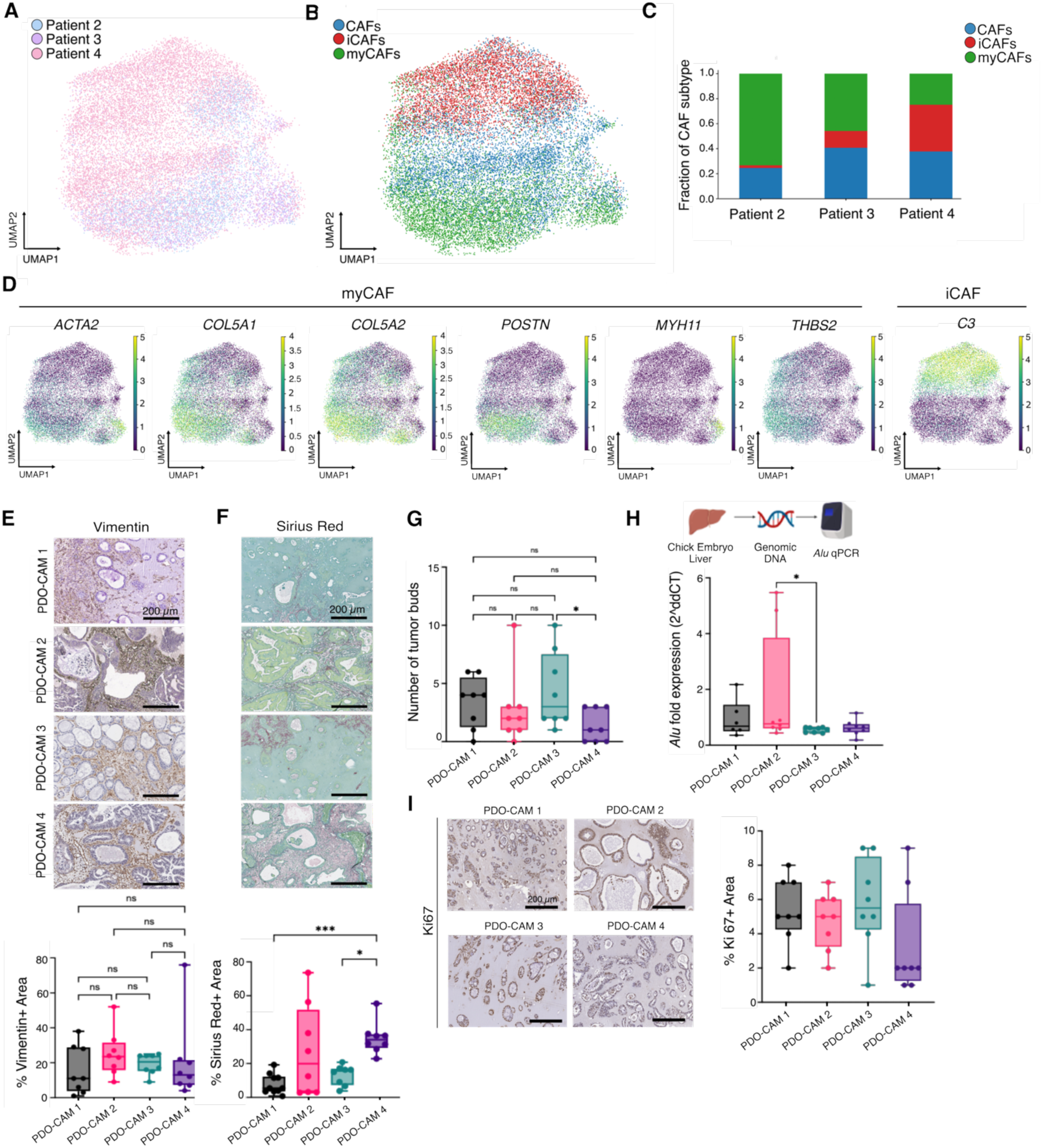
Patient tumor CAF heterogeneity informs the modeling of tumor-stroma interactions and proliferation-independent dissemination *in ovo*. (A) UMAP visualization of CAFs derived from Xenium spatial transcriptomic profiling across patient tumors. (B) UMAP projection annotating distinct patient CAF sub-populations, highlighting iCAFs, myCAFs, and an unaligned CAF population displaying intermediate marker phenotypes. (C) Compositional bar plot quantifying the relative proportions of CAF subpopulations across individual patient tumors. (D) UMAPs depicting the expression of myCAF markers (*ACTA2*, *COL5A1*, *COL5A2*, *POSTN*, *MYH11*, *THBS2*) and the iCAF marker (*C3*) in the patient tumors. (E) Representative immunohistochemical staining (at 20X magnification) and quantification of host-derived chick fibroblasts using a chicken-specific Vimentin antibody across PDO-CAM tumors. Scale bar = 200 μm. (F) Representative immunohistochemistry images (at 20X magnification) and quantitative analysis of Sirius Red staining of PDO-CAM tumors. Scale bar = 200 μm. (***P = 0.0003; *P=0.0183). (G) Quantification of malignant tumor budding at the invasive peripheral fronts of the PDO-CAM tumors. (*P=0.0310). (H) *Alu* qPCR of chick embryo liver to measure metastatic dissemination (*P=0.0332). (I) Representative immunohistochemistry (at 20X magnification) and quantitative analysis of Ki67 expression within PDO-CAM tumors. Scale bar = 200 μm. For all *in ovo* PDO-CAM experiments (E–I), sample sizes are *n* = 8 per PDO line. Statistical significance determined by Kruskal-Wallis test. Data are presented as median with Min to Max range; ns = non-significant.

The prominent desmoplastic features observed in patient tumours, including stromal infiltration and ECM deposition, suggested active communication between tumor cells and the surrounding microenvironment. We therefore exposed naïve human pancreatic stellate cells (hPSCs) to CM derived from the four PDO lines and assessed hPSC activation. Upon exposure to PDO-CM, hPSCs, similar to mPSCs, underwent a dramatic morphological transformation, transitioning from a quiescent state into an elongated, spindle-shaped morphology with prominent α-SMA expression (Supplementary Figure 7C, Supplementary Figure 4B). qPCR validation of these activated cells confirmed a robust up-regulation of *ACTA2* and *COL1A1* transcripts, driven most prominently by supernatants from PDO 2 and PDO 4 (Supplementary Figure 7D-E).

To determine if this tumor-driven activation successfully recapitulates the patient stromal architecture within our *in ovo* model, we analyzed the stromal composition of the PDO-CAM tumors. As seen in the KPC-CAM model, PDOs transplanted onto the CAM successfully activated the naïve CAM microenvironment and recruited α-SMA-positive stromal cells into the tumor periphery (Supplementary Figure 8A). These findings further mirrored the KPC-on-CAM model with immunohistochemistry for chicken-specific Vimentin demonstrating an extensive infiltration of host avian fibroblasts and encapsulating the human tumor areas, confirming robust recruitment into the TME by PDOs (Figure 6E). Mirroring our *in vitro* activation findings, PDO-CAM 2 and PDO-CAM 4 displayed the highest percentage of Sirius Red-positive area *in ovo*, demonstrating a preserved link between tumor secretome potency and structural desmoplasia (Figure 6F). Multiplex immunofluorescence co-staining for chicken-specific Vimentin, IL-6, and HABP further confirmed that the *in ovo* model accurately organizes localized CAF signaling alongside heavy hyaluronan deposition; notably, PDO-CAM 1 and PDO-CAM 3 significantly accelerated this matrix deposition (Supplementary Figure 8B-C).

To evaluate the functional consequences of this tumor-stroma crosstalk, we investigated the metastatic potential of the cells. *In vitro* scratch assays revealed variable intrinsic migratory capacities among the lines, with 2D lines generated from PDO 3 and PDO 4 executing accelerated, significant wound closure by 24 hours, while lines PDO 1 and PDO 2 remained comparatively slower (Supplementary Figure 8D-E). Similarly, Matrigel invasion assays demonstrated that the addition of CAFs (+CAF) or CAM tissue fragments (+CAM) to the lower chamber acted as a potent chemoattractant, significantly and comparably boosting the invasive and metastatic potential of PDO 3 and PDO 4 (Supplementary Figure 8F). This finding is highly consistent with established paradigms showing that CAFs broadly enhance the invasive capacity and EMT-like programs of pancreatic epithelial cells (Sahai et al., 2020).

In alignment with these findings, histopathological analysis of the *in ovo* tumors revealed prominent malignant tumor budding at the invasive peripheral fronts, with PDO 3 showing the highest number of tumor buds (Figure 6G). To track systemic dissemination, we performed highly sensitive human-specific *Alu* qPCR on host chick embryo livers, confirming early metastatic dissemination to the distant organ site, with PDO-CAM 2 exhibiting a high-burden dissemination profile (Figure 6H). Finally, to determine whether this metastatic spread was driven by accelerated cell division or an intrinsic invasive phenotype, we quantified cell proliferation via Ki-67 staining (Figure 6I). Proliferation rates did not correlate with metastatic burden; despite its high metastatic dissemination, PDO-CAM 2 showed no difference in proliferation compared to other lines. This demonstrates that early systemic dissemination in this model occurs as a function of active, CAF-mediated invasion regardless of primary cellular proliferation rates.

## Discussion

PDAC is characterized by continuous reciprocal interactions between malignant cells and a highly desmoplastic TME. Rather than evolving independently, tumor and stromal compartments undergo coordinated adaptation, with tumor cell states influencing stromal composition and function while stromal signals shape tumor cell plasticity, subtype identity, and invasive behavior (Klein et al., 2025). Understanding these processes of tumor–stroma co-evolution is therefore essential for elucidating the mechanisms underlying PDAC progression, therapeutic resistance, and metastatic dissemination. However, experimental models that preserve both tumor heterogeneity and stromal complexity remain limited. In this study, we establish KPC-on-CAM and PDO-on-CAM platforms as rapid, scalable *in ovo* models that enable the reconstruction of tumor architecture, stromal remodeling, and the emergence of heterogeneous tumor and stromal populations. Through integrated sc-RNAseq, histological, and spatial transcriptomic analyses, we demonstrate that the CAM microenvironment supports key processes of tumor–stroma co-evolution that are largely lost under *in vitro* conditions.

A central observation of this study is the profound reduction in tumor cell heterogeneity following adaptation of KPC tumors to standard 2D culture conditions. scRNA-seq analyses revealed that endogenous KPC tumors contained multiple transcriptionally distinct tumor cell populations encompassing diverse subtype-associated programs, whereas corresponding cell lines displayed marked phenotypic convergence and loss of cellular heterogeneity. These findings are consistent with previous studies demonstrating that conventional *in vitro* culture impose strong selective pressures that favor the expansion of a limited number of proliferative cell states while eliminating microenvironment-dependent phenotypes (Khaliq et al., 2024). This phenotypic convergence not only diminishes the cellular complexity of the tumor but also disrupts the dynamic reciprocal interactions between malignant cells and the surrounding stroma that are fundamental drivers of PDAC progression. These observations highlight the limitations of traditional *in vitro* systems for modeling the dynamic processes governing tumor–stroma co-evolution.

Remarkably, transplantation of KPC cells onto the CAM resulted in the re-establishment of parental tumor architecture. Histological analyses demonstrated restoration of ductal morphology, tissue compartmentalization, and invasion-associated growth patterns resembling those observed in endogenous tumors. These findings suggest that tumor architecture is not solely encoded by intrinsic genetic programs but can be reconstituted when tumor cells are exposed to a permissive microenvironment. Increasing evidence indicates that architectural states reflect biologically meaningful programs associated with differentiation, invasion, and disease progression (Di Chiaro et al., 2024). The ability of KPC cells to reconstruct tumor architecture within the CAM microenvironment therefore supports the concept that environmental cues play a critical role in regulating tumor organization and cellular identity.

Importantly, restoration of tumor architecture was accompanied by active remodeling of the surrounding microenvironment. KPC-on-CAM tumors induced ECM deposition, stromal compartmentalization, and the emergence of CAF-rich regions that closely resembled key features of the pancreatic TME. These observations indicate that tumor cells are not passive occupants of a pre-existing niche but active architects of their microenvironment. The ability of KPC cells to recruit and organize stromal elements within a developmentally distinct host tissue highlights the robustness of tumor-intrinsic programs that drive microenvironmental remodeling. Notably, the emergence of stromal features occurred in parallel with the restoration of complex tissue architecture, suggesting that tumor organization and stromal evolution are interconnected processes that develop simultaneously. Together, these findings provide direct evidence that the CAM platform can capture fundamental aspects of tumor-driven stromal evolution that characterize PDAC progression.

Extending these observations to the human system, PDO-on-CAM tumors faithfully recapitulated patient-specific architectural features of the original tumors, underscoring the translational relevance of the model. By preserving structural characteristics that reflect underlying tumor biology, the platform bridges the gap between reductionist organoid cultures and more complex *in vivo* models. The ability of PDOs to reconstruct these patient-specific tumor architectures suggests that critical developmental and microenvironmental programs remain preserved during organoid culture and can be reactivated following engraftment into a permissive *in vivo* niche. PDO-on-CAM tumors capture clinically meaningful aspects of disease heterogeneity, providing a rapid and scalable system for patient-specific functional studies, therapeutic response assessment, and biomarker development.

Spatial transcriptomic profiling further demonstrated that PDO-CAM tumors preserve key aspects of tumor cell heterogeneity and subtype architecture observed in the parental patient tumors. Despite exhibiting reduced overall cellular complexity relative to clinical specimens, PDO-CAM tumors retained heterogeneous tumor cell states and maintained subtype-associated transcriptional programs. Distance-to-meta-cell analyses confirmed that much of the transcriptional diversity present in patient tumors was conserved following engraftment. These findings contrast sharply with the phenotypic convergence observed in conventional *in vitro* models and suggest that preservation of tumor heterogeneity is facilitated by the establishment of a supportive stromal niche. Rather than serving merely as a structural scaffold, the reconstructed microenvironment appears to provide signals that sustain cellular diversity and subtype-specific programs. This capacity to maintain both architectural and transcriptional complexity underscores the model’s utility for investigating tumor plasticity, subtype evolution, and mechanisms governing intratumoral heterogeneity.

CAF heterogeneity has emerged as a critical determinant of PDAC progression, influencing ECM remodeling, immune regulation, therapeutic resistance, and metastatic dissemination (Oh et al., 2023). Spatial transcriptomic analyses of patient tumors identified distinct CAF populations, consistent with the well-established stromal heterogeneity that characterizes PDAC. Stromal features consistent with CAF differentiation were also observed in PDO-on-CAM tumors by immunohistochemical analyses, indicating that PDOs retain the capacity to induce and organize stromal populations within the CAM microenvironment. Although the transcriptional identity of CAF subsets was not directly resolved in the PDO-on-CAM setting, the emergence of CAF-associated stromal compartments suggests that key tumor-derived signaling pathways involved in stromal recruitment and activation remain operational following engraftment. Conversely, the persistence of heterogeneous tumor cell states alongside diverse CAF populations suggests that reciprocal signaling networks between these compartments are preserved. The conservation of heterogeneity in both tumor and stromal compartments provides compelling evidence that the model captures ongoing processes of tumor–stroma co-evolution.

Consistent with the established relationship between tumor cell state and stromal remodeling, we observed differences in ECM composition and organization. Variations in collagen and hyaluronan deposition, cytokine signaling networks, and overall stromal architecture were evident between tumors, suggesting that distinct tumor cell populations actively shape their surrounding microenvironment. These findings further support the concept that tumor and stromal compartments evolve together through continuous reciprocal interactions (Niu et al., 2024). The ability to functionally interrogate these stromal phenotypes *in ovo* offers a valuable opportunity to dissect mechanisms through which tumor cell identity influences disease aggressiveness and therapeutic response.

Another important strength of the platform is its capacity to quantify early metastatic dissemination. Using species-specific qPCR targeting *B1* and *Alu* DNA elements in murine and human cells, respectively, we achieved sensitive detection of disseminated tumor cells within embryonic tissues. This provides a robust, quantitative measure of metastatic potential and enables investigation of dissemination events that are difficult to capture in conventional preclinical models. Importantly, integration of metastatic assessment with analyses of tumor architecture, stromal remodeling, and cellular heterogeneity creates a comprehensive framework for studying how tumor–stroma interactions contribute to metastatic progression.

From a translational perspective, the PDO-on-CAM platform offers substantial opportunities for functional stratification of PDAC. The preservation of subtype-specific stromal remodeling, CAF heterogeneity, and metastatic behavior suggests that patient-derived tumors can be classified not only by their molecular characteristics but also by their capacity to shape and respond to the surrounding microenvironment. Such functional phenotyping may facilitate identification of clinically relevant biomarkers and therapeutic vulnerabilities that are not apparent from tumor-intrinsic analyses alone.

The practical advantages of the CAM model further enhance its translational utility. Compared with murine models, the CAM platform is inexpensive, scalable, technically accessible, and characterized by a rapid experimental turnaround. These features make it particularly attractive for medium-throughput drug screening, biomarker discovery, genetic perturbation studies, and personalized oncology applications where timely generation of functional data is essential. While conventional mouse models remain indispensable for studying long-term tumor evolution and immune interactions, the CAM platform provides a complementary approach that substantially accelerates mechanistic and translational investigations.

In summary, we demonstrate that the CAM microenvironment supports reconstruction of the reciprocal interactions that define PDAC biology. Both KPC-on-CAM and PDO-on-CAM models allow tumor cells to actively remodel their surroundings to generate a PDAC-like stroma, preserving parental tumor architecture, tumor cell heterogeneity, subtype identity, and CAF diversity, which are lost in conventional *in vitro* systems. These findings establish the *in ovo* CAM model as a powerful and versatile platform for investigating the dynamic processes through which tumor and stromal populations jointly evolve during pancreatic cancer progression and for advancing translational research in PDAC.

## Acknowledgments

We thank all members of the Reichert laboratory for their valuable scientific input and insightful discussions throughout this study. We are grateful to Lisa Fricke for clinical study support and assistance with patient enrolment in the PDAC cohort. We further thank Thomas Metzler, Inken Stahl, Tanja Groll, Olga Seelbach, Petra Hauke, Marco Heinrichs, Nadja Lewandowski-Hoppe, and Leona Arps for their excellent technical assistance.

We gratefully acknowledge the following funding sources: M.R. was supported by Deutsche Krebshilfe through the Max Eder Program (grants 111273 and 70114328) and by the DFG (grants RE 3723/4-1, RE 3723/6-1, and DFG–SFB1321, Project ID 329628492, Project 12). Further support for M.R. was provided by the Biosystems Design Munich (BioSysteM): Multicellular Systems and Organoids – DFG Cluster of Excellence; the German Federal Ministry of Education and Research (BMBF) through the projects SATURN3 (01KD2206P), QuE-MRT (13N16450), and FAIrPaCT2 (01KD2414C); the German Cancer Consortium (DKTK) Strategic Initiative Organoid Platform; and the Bavarian Center for Cancer Research (BZKF) Lighthouse Project “Preclinical Model Systems.” A.R. was supported by the Juan Rodés PhD Student Fellowship from the European Association for the Study of the Liver (EASLJR2002-01).

Figure illustrations were created using BioRender.com; all associated licenses were obtained by M.R.

