## Supplementary Figures and Legends for "Modeling pancreatic cancer tumor stroma co-evolution in an *in ovo* model"

Supplementary Figure 1

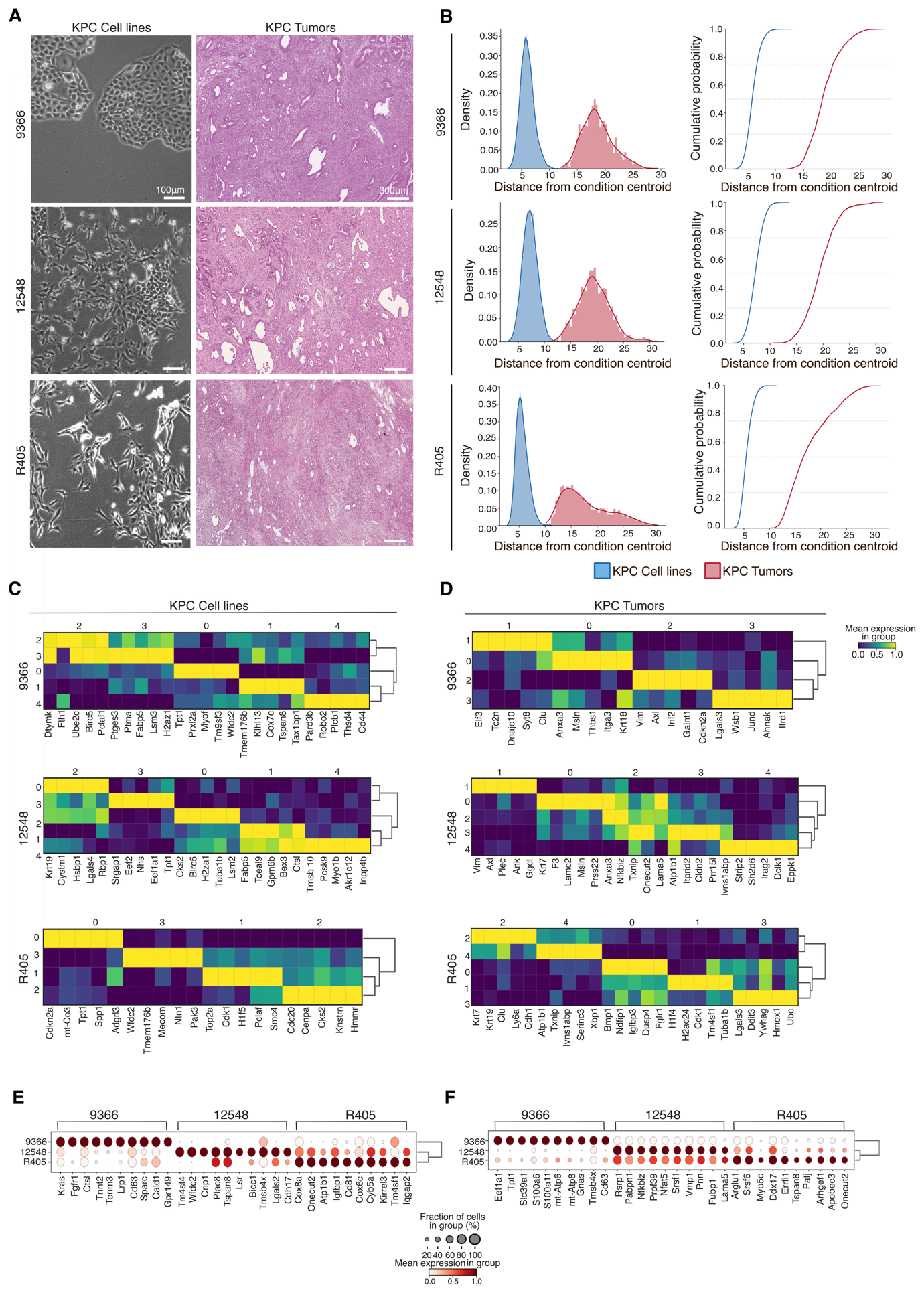

Supplementary Figure 2

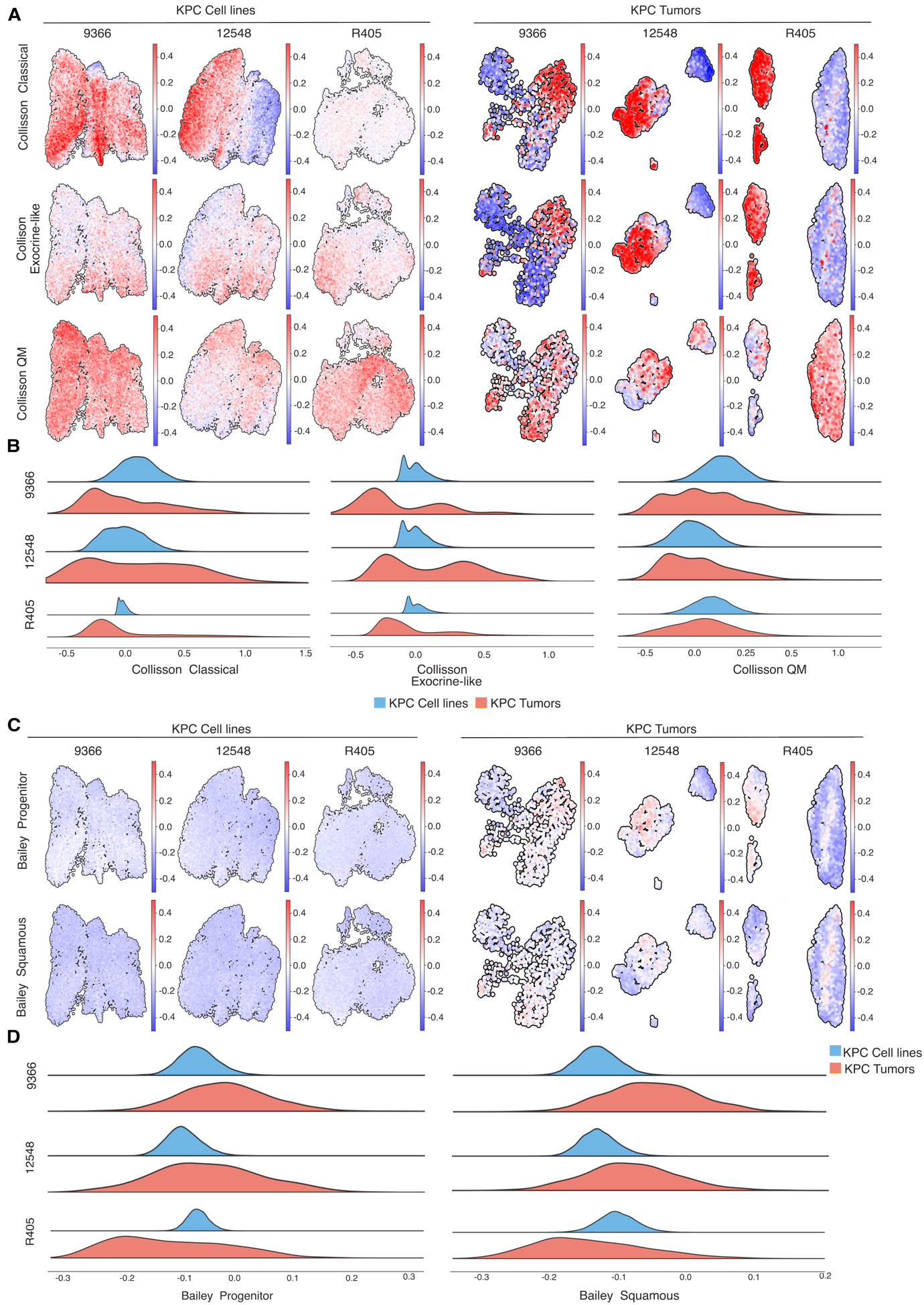

Supplementary Figure 3

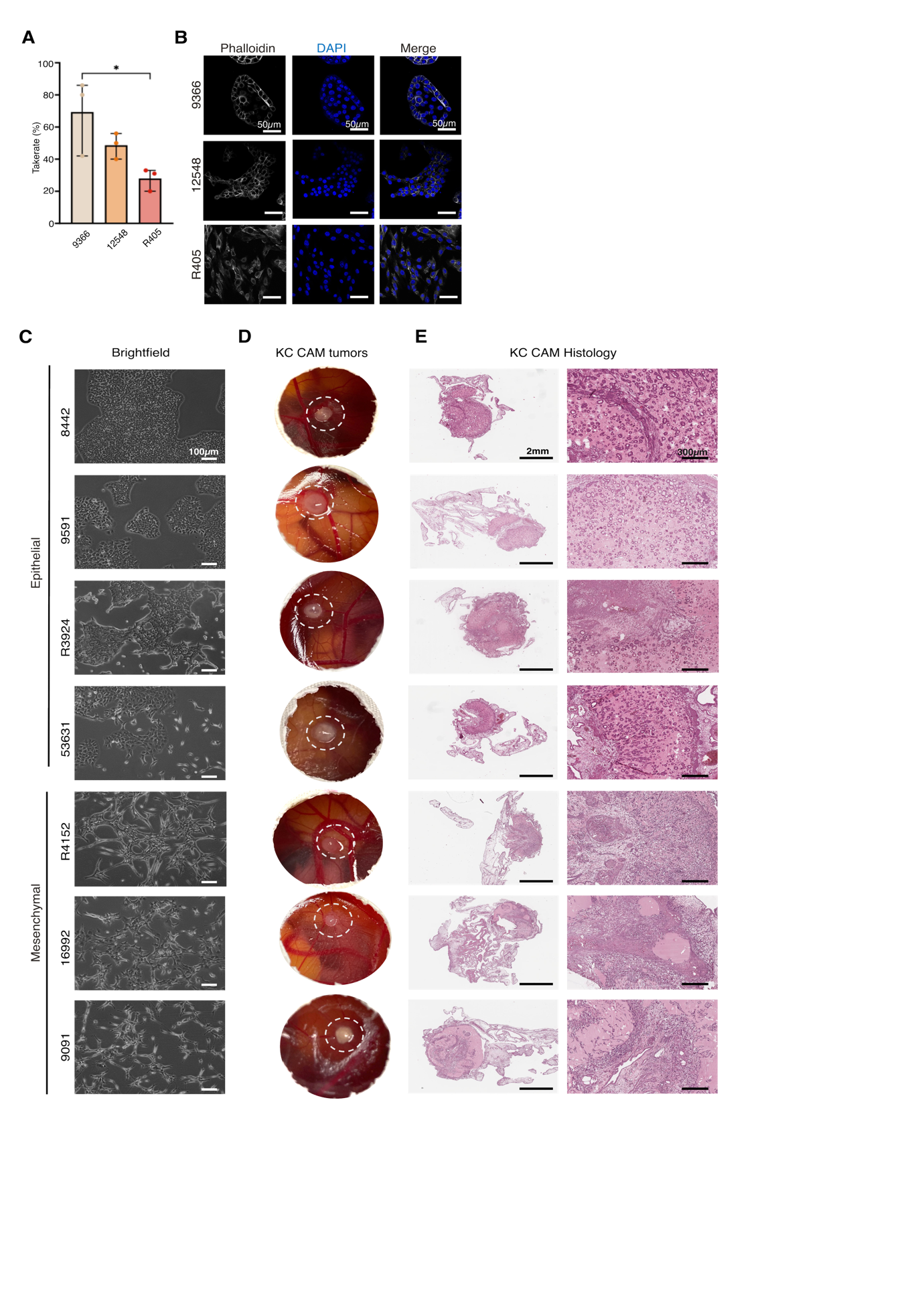

Supplementary Figure 4

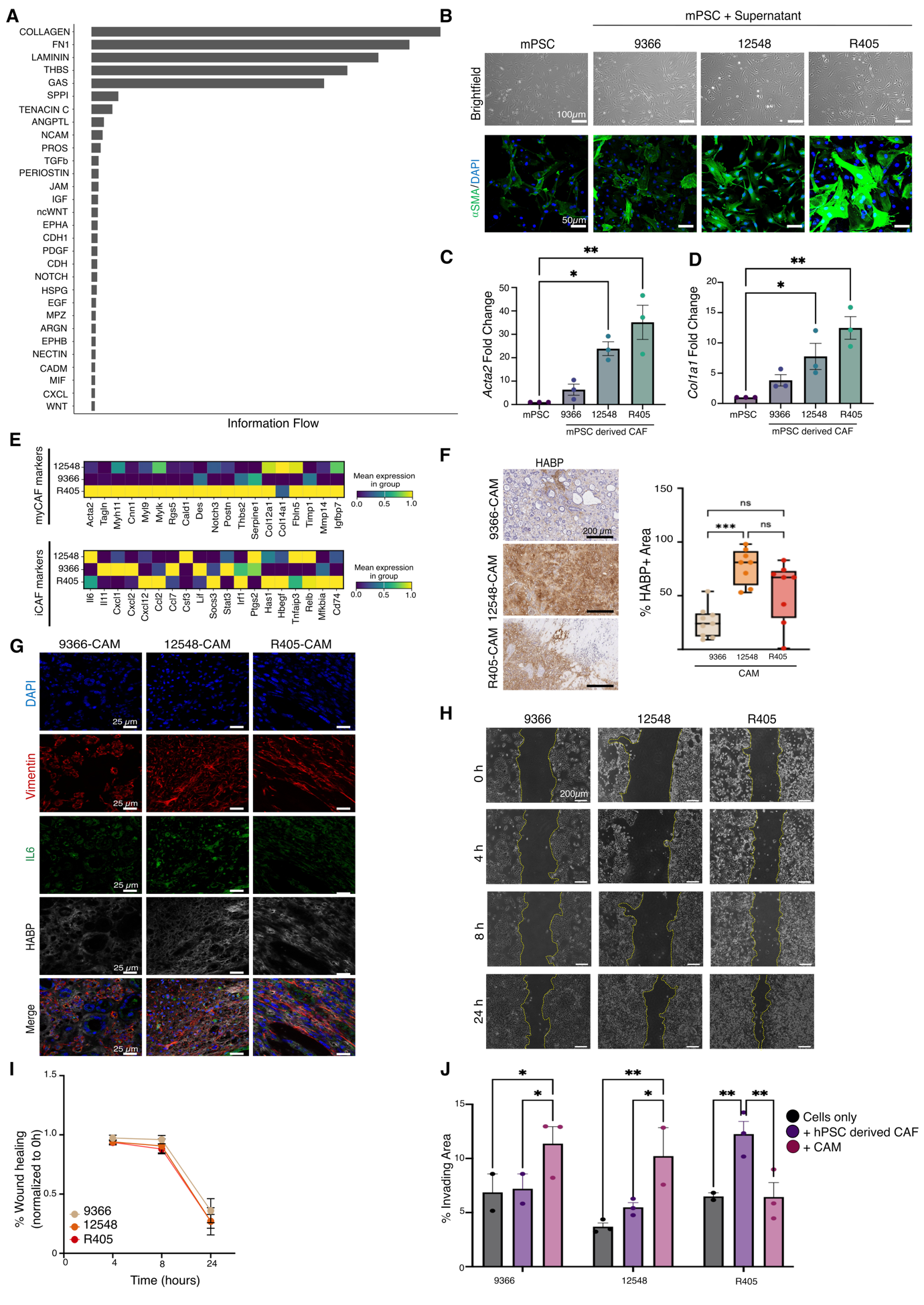

Supplementary Figure 5

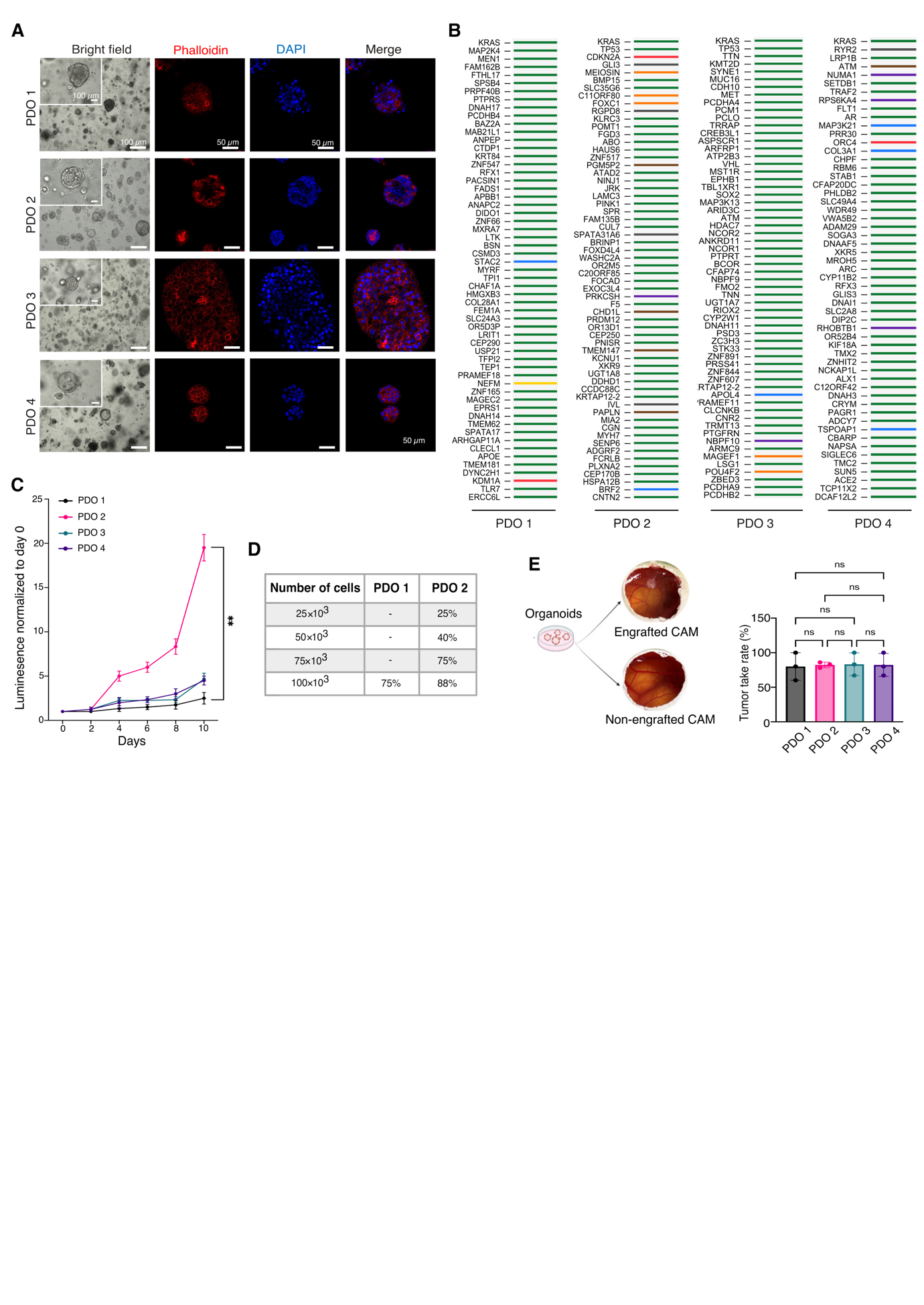

Supplementary Figure 6

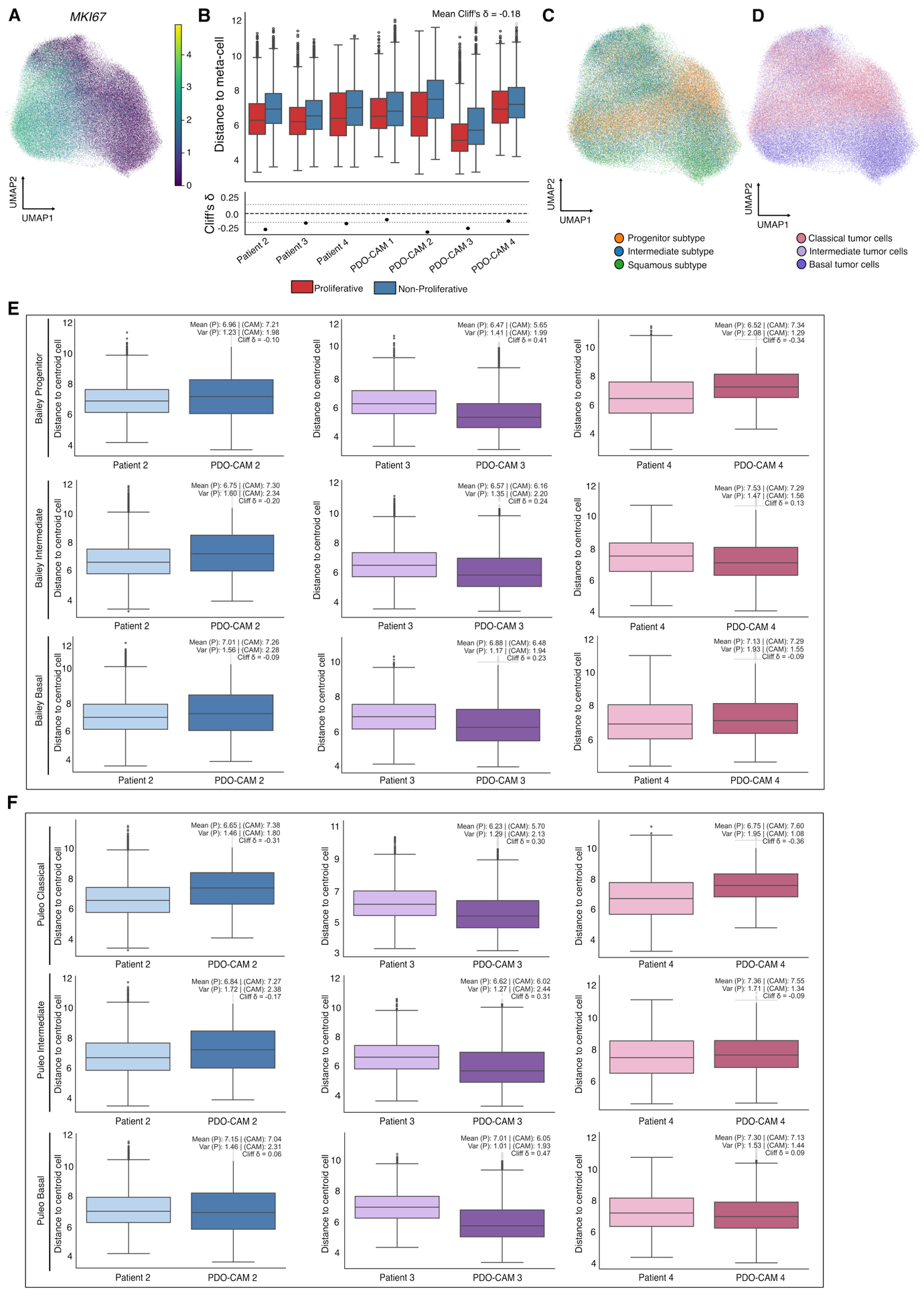

Supplementary Figure 7

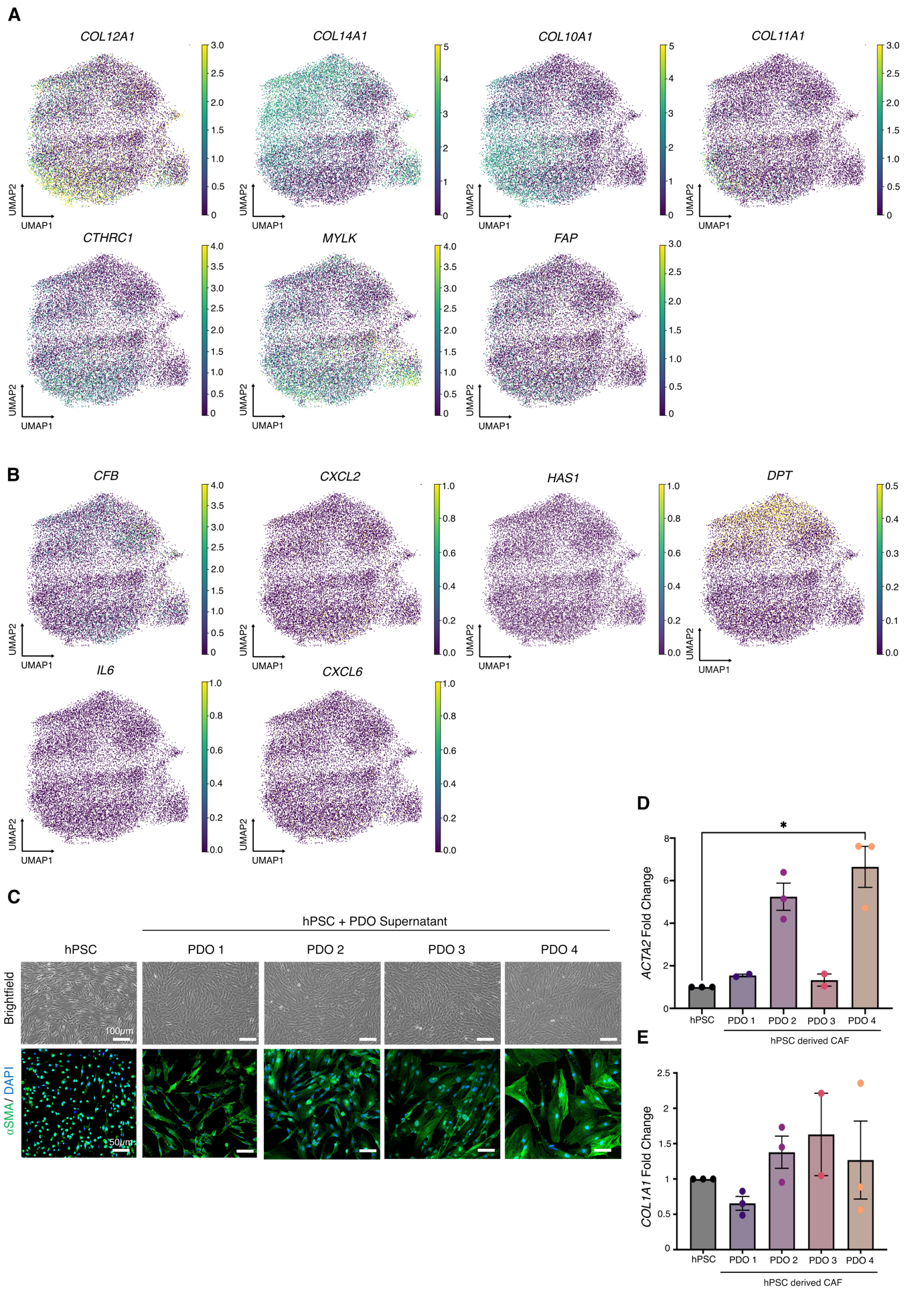

Supplementary Figure 8

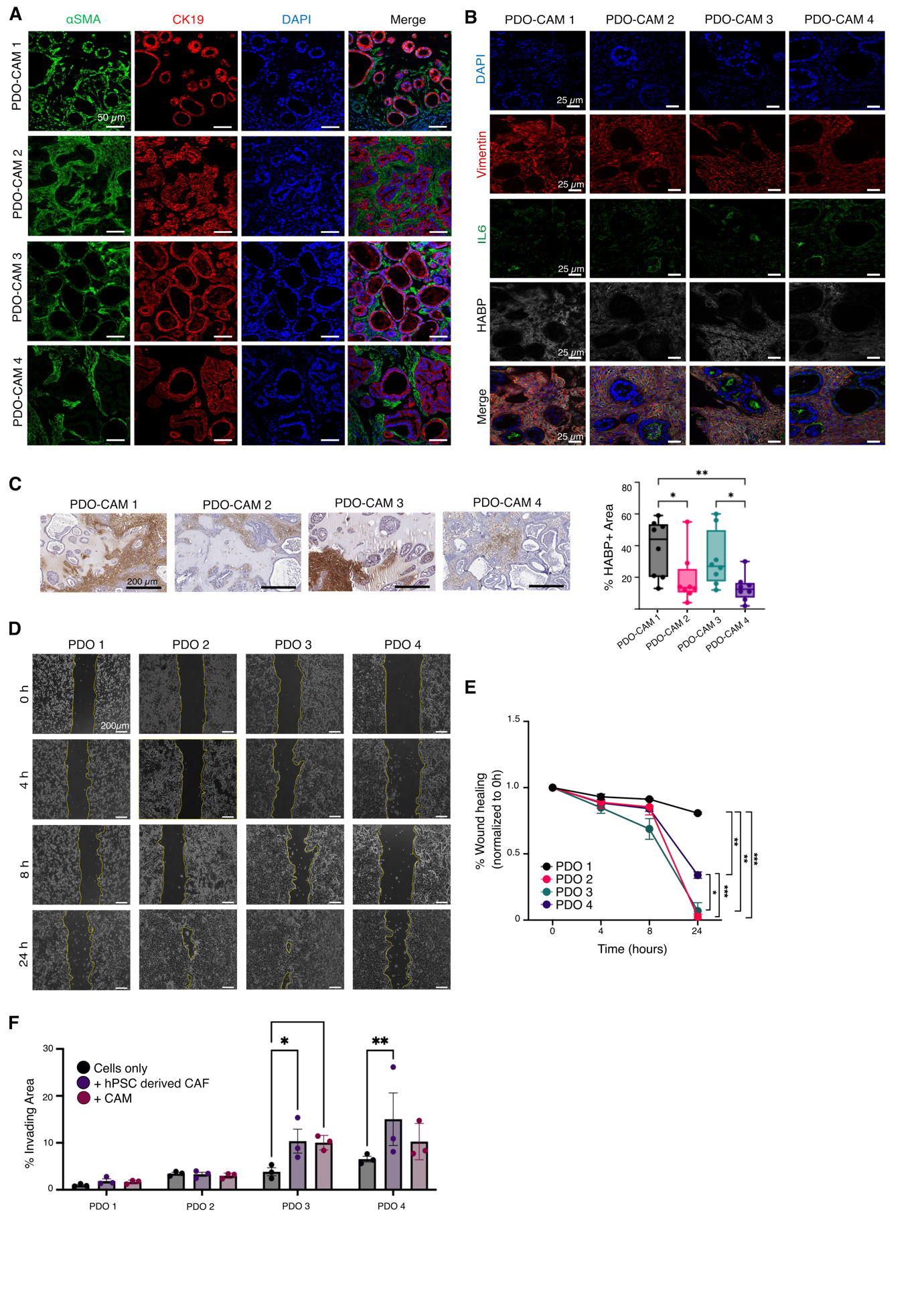

**Supplementary Figure Legends**

Supplementary Figure 1. Extended Analyses of Cellular States and Cluster-Specific Gene Expression in KPC cell lines and parental KPC tumors.

**(A)** Representative bright-field images (at 10X magnification) of KPC cell lines (left, scale bar = 100 µm) and corresponding H&E sections of parental KPC tumors (at 8X magnification) (right, scale bar = 300 µm), illustrating morphological features *in vitro* and *in vivo.* **(B)** Density plots of cell-to-centroid distances (left) and cumulative probability plots (right) for KPC cell lines and KPC tumors. **(C)** Heatmap of Leiden cluster marker genes identified from sc-RNAseq data of KPC cell lines, highlighting cluster-specific transcriptional programs. **(D)** Heatmap of Leiden cluster marker genes identified from matched parental tumors, revealing transcriptional diversity and tumor-specific cellular states. **(E)** Dot plot showing selected differentially expressed genes identified through comparative analysis between the KPC cell lines. **(F)** Dot plot showing selected differentially expressed genes identified through comparative analysis between the KPC tumors.

Supplementary Figure 2. Collisson and Bailey subtype distributions across KPC cell lines and parental KPC tumors

**(A)** UMAP projections of sc-RNAseq data of tumor cells from KPC cell lines and corresponding parental tumors, annotated according to **Collisson PDAC subtype scores**. **(B)** Ridge plots showing the distribution of Collisson subtype scores across tumor cells from each KPC cell line and parental KPC tumor sample, highlighting differences in subtype composition and transcriptional heterogeneity. **(C)** UMAP projections annotated by **Bailey PDAC subtype scores** of tumor cells from KPC cell lines and parental KPC tumors. **(D)** Ridge plots showing the distribution of Bailey subtype scores across tumor cells from each KPC cell line and parental KPC tumor sample.

Supplementary Figure 3. **Morphological and Histological Characterization of KC Murine Cell Lines and Derived CAM Tumors**.

(A) Tumor take-rate for KPC-CAM. Data represent cumulative successful grafts out of total embryos engrafted across 3 independent experimental runs with 6-8 eggs/experiment. Statistical significance was determined by Kruskal-Wallis test (*p=0.0113). (B) Immunofluorescence for Phalloidin (gray) and DAPI (blue) (at 40X magnification) of KPC-cell lines showing the epithelial and mesenchymal morphology. Scale bar = 50 µm (C) Brightfield microscopy (at 10X magnification) of KC cell lines of epithelial (8442, 9591, R3924, 53631) and mesenchymal (R4152, 16992, 9091) phenotypes. Scale bar = 100 µm. (D) Macroscopic images of KC-CAM tumors. (E) H&E of KC-CAM tumors (left; at 2X magnification) Scale bar = 2mm. Right (at 10X magnification). Scale bar = 300 μm.

Supplementary Figure 4. Extracellular matrix crosstalk networks and functional tumor-mediated activation of mouse pancreatic stellate cells.

(A) RankNet plot ranking the relative information flow of signalling pathways within the TME, showing the top 30 upregulated signalling networks. (B) Representative brightfield field images (top, at 10X magnification, scale bar = 100 μm) and immunofluorescence staining images (bottom, at 20X magnification, scale bar = 50 μm) showing the activation and morphology of mPSCs stained for $⍺$-SMA (green)/DAPI (blue) after incubation with conditioned media from KPC cell lines. (C-D) qPCR showing fold changes in mRNA transcript expression of (C) *Acta2* and (D) *Col1a1* in mPSC derived CAFs. Data presented as mean with SEM. Statistical significance determined by Kruskal-Wallis (*n*=3). (C) *p=0.0226; **p=0.0044 and (D) *p=0.0401; **p=0.0030. (E) Heatmaps showing relative transcriptional expression levels of established myCAF markers (top) and iCAF markers (bottom) in KPC tumors. (F) Representative immunohistochemistry (at 20X magnification) and corresponding quantitative bar plots measuring the percentage of HABP+ areas. Scale bar = 200 μm. Sample sizes are *N*=9for 9366 KPC-CAM, *N*=8 for 12548 KPC-CAM, and *N*=9 for R405 KPC-CAM tumors. Statistical significance was determined by the Kruskal-Wallis test (***p=0.0006). Data are presented as median with Min to Max range; ns = non-significant. (G) Multiplex immunofluorescence images (at 63X magnification) showing expression of DAPI (blue) Vimentin (red), IL-6 (green), and HABP (gray) in KPC-CAM xenografts. Scale bar = 25 μm. (H) Representative phase contrast time-course imaging (at 5X magnification) and (I) quantification of scratch-wound closure across individual KPC lines. Scale bar = 200 μm. The data were collected from 3 independent experiments (*n*=3). Statistical significance determined by 2way Analysis of Variance (ANOVA). ns = non-significant. Data are presented as mean and SEM. (J) Invasion analysis showing percentage of invading cells through a Matrigel coated membrane when stimulated by CAFs (+CAF) and CAM tissue (+CAM). Statistical significance determined by 2way Analysis of Variance (ANOVA) (9366*p=0.0314, 0.0438; 12548 **p=0.0.0038, *p=0.0.0247; R405 **p=0.0039, 0.0085). The data were collected from 3 independent experiments (*n*=3). Data are shown as mean with SEM.

Supplementary Figure 5. **Clinical, Molecular, and Functional Characterization of PDOs**

(A) Mutation profiles of the PDOs, showing the top 58 mutations by allelic frequency. (B) Growth rates of the PDO lines over 10 days by CellTiter-Glo® assay. Statistical significance was determined by Friedman test (**p=0.0017). Data are shown as mean with SEM. (C) Optimization of PDO seeding density on CAM using the PDO lines with the highest and lowest *in vitro* proliferation, PDO 2 and PDO 1, respectively. At least 1×10^5^ cells were required to achieve a 75% tumor take rate for the slowest-growing PDO line, PDO 1. (D) PDO take-rate on the CAM. Data represent cumulative successful grafts out of total embryos engrafted across 3 independent experimental runs with 6-8 eggs/experiment. ns = non-significant

Supplementary Figure 6. Preservation of the Proliferative Landscape and Extended Transcriptomic Subtypes

(A) UMAP projection of *MKI*67 expression across the tumor cell population (B) Box plot showing transcriptomic heterogeneity across proliferative and non-proliferative tumor cell populations across the samples. **(C)** UMAP of tumor cells classified according to Bailey molecular subtypes (Progenitor, Intermediate, and Squamous) across both patient and PDO-CAM tumors. **(D)** UMAP of tumor cells classified according to Puleo molecular subtypes (Classical, Intermediate, and Basal) across both patient and PDO-CAM tumors. (E**)** Box plots showing subtype heterogeneity according to Bailey subtyping. (F**)** Box plots showing subtype heterogeneity according to Puleo subtyping.

Supplementary Figure 7. Extended spatial characterization of CAF subpopulation markers and tumor mediated *in vitro* stellate cell activation.

(A) Supplementary Xenium UMAPs displaying the expression profile of additional myCAF markers (*COL12A1*, *COL14A1*, *COL10A1*, *COL11A1*, *CTHRC1*, *MYLK*, and *FAP*) across the patient tumors. (B) UMAPs showing the expression profiles of additional iCAF markers (*CFB*, *CXCL2*, *HAS1*, *DPT*, *IL6*, and *CXCL6*). (C) Representative brightfield images (top, at 10X magnification, scale bar = 100 μm) and immunofluorescence (bottom, at 20X magnification, scale bar = 50 μm) showing α-SMA (green) and DAPI (blue) of hPSCs following incubation with CM from PDOs. (D–E) mRNA expression levels quantified via qPCR for (D) *ACTA2* and (E) *COL1A1* in hPSCs following activation with PDO supernatant. Data are presented as mean with SEM. Significance determined by Kruskal-Wallis test (*p=0.0316).

Supplementary Figure 8. Stromal characterization of PDO-CAM tumours and functional influence of stromal cells on PDO migration.

(A) Representative immunofluorescence staining (at 40X magnification) of ⍺-SMA (green) and CK19 (red) showing stromal infiltrates in PDO-CAM tumors. Scale bar = 50 μm. (B) Multiplex immunofluorescence images (at 63X magnification) displaying co-localization of DAPI (blue), chicken-specific Vimentin (red), IL-6 (green), and HABP (gray) in PDO-CAM tumors. Scale bar = 25 μm. (C) Representative immunohistochemical images (at 20X magnification) and matching quantification of HABP+ areas in PDO-CAM tumors. Scale bar = 200 μm. Data are presented as median with Min to Max range (N=8 per PDO line). Statistical significance determined by Kruskal-Wallis test (*p=0.0183, 0.0260; **p=0.0068). (D) Representative phase contrast time-course imaging (at 5X magnification) and (E) quantification of scratch-wound closure across PDO lines. Scale bar = 200 μm. The data were collected from 3 independent experiments (*n*=3). Statistical significance determined by 2way ANOVA. ns=non-significant. *p=0.0362, **p=0.0012, 0.0065, ***p=0.0001, 0.0007. Data are represented as mean with SEM. (F) Matrigel invasion assay quantification demonstrating significantly elevated invasion of tumor cells driven by CAFs (+CAF) and CAM (+CAM). Statistical significance determined by ANOVA (9366*p=0.0.0314, 0.0438; 12548 **p=0.0.0038, *p=0.0.0247; R405 **p=0.0039, 0.0085). (PDO 3 *p=0.0333, 0.0257; PDO 4 *p=0.0050). The data were collected from 3 independent experiments (*n*=3). Data are shown as mean with SEM.

**Supplementary Table Legends**

Supplementary Table 1. Patient data associated with the PDOs used in this study.

Supplementary Table 2. Xenium panel comprising 377 pre-designed and 100 custom genes.
